# Spectral-switching analysis reveals real-time neuronal network representations of concurrent spontaneous naturalistic behaviors in human brain

**DOI:** 10.1101/2024.07.08.600416

**Authors:** Hongkun Zhu, Andrew J. Michalak, Edward M. Merricks, Alexander H. C. W. Agopyan-Miu, Joshua Jacobs, Marla J. Hamberger, Sameer A. Sheth, Guy M. McKhann, Neil Feldstein, Catherine A. Schevon, Elizabeth M. C. Hillman

## Abstract

Despite abundant evidence of functional networks in the human brain, their neuronal underpinnings, and relationships to real-time behavior have been challenging to resolve. Analyzing brain-wide intracranial-EEG recordings with video monitoring, acquired in awake subjects during clinical epilepsy evaluation, we discovered the tendency of each brain region to switch back and forth between 2 distinct power spectral densities (PSDs 2-55Hz). We further recognized that this ‘spectral switching’ occurs synchronously between distant sites, even between regions with differing baseline PSDs, revealing long-range functional networks that would be obscured in analysis of individual frequency bands. Moreover, the real-time PSD-switching dynamics of specific networks exhibited striking alignment with activities such as conversation and hand movements, revealing a multi-threaded functional network representation of concurrent naturalistic behaviors. Network structures and their relationships to behaviors were stable across days, but were altered during N3 sleep. Our results provide a new framework for understanding real-time, brain-wide neural-network dynamics.

## Introduction

Over 30 years of functional imaging studies have demonstrated that the human brain operates as a complex and interconnected system, with distinct functional networks and long-range coordination of neural activity^1^. It is also well-recognized that cognitive and behavioral functions are not executed by individual brain regions in isolation; but require collaboration between multiple distinct brain regions^2,3^. Yet, how our brains coordinate our behavior from moment to moment, permitting us to think, talk and move at the same time, has been almost impossible to decode.

Functional magnetic resonance imaging (fMRI) studies have demonstrated the presence of widespread brain networks by analyzing cross-areal correlations in the blood oxygen level-dependent (BOLD) signal^1,4,5^. However, interpretability is debated, as the BOLD signal reflects an indirect measurement of neural activity^6^. Constraints on fMRI data acquisition also necessitate the collection of behaviorally-restricted, and relatively short duration ‘resting state’ datasets which cannot capture longer term patterns of dynamic coordination^7,8^ and their relationship to naturalistic behaviors. Electroencephalography (EEG) and intracranial electrocorticography (iEEG) capture dynamic neural activity as the summation of postsynaptic currents. Scalp EEG^9–11^, iEEG^12–18^ and magnetoelectroencephalography (MEG)^19–21^, have been used to characterize long-range coordination and network patterns, in many cases corroborating fMRI observations of ‘functional connectivity’^22^, measurements. However, EEG and iEEG generally have limited spatial sampling, presenting challenges for investigation of large-scale interactions across multiple brain regions, while interpretation has also been confounded by discrepancies in frequency band selection across prior studies^23–25^. As a result, few studies have been able to harmonize functional connectivity observations across modalities and link their activity patterns to real-time behavior.

The invasive, long-term, and often multi-regional iEEG monitoring utilized for epilepsy surgery evaluation presents a valuable opportunity for studying brain-wide dynamic neural activity in behaving human subjects. In this study, we analyzed over 93 hours of iEEG recordings along with simultaneously acquired video recordings from 10 patients with drug-resistant focal epilepsy syndromes, who underwent invasive iEEG with broadly distributed bilateral depth electrodes for clinical evaluation.

Rather than representing data as time-varying signals within specific frequency bands, we incorporated all frequencies (2-55 Hz) into our analysis by calculating the power spectral density (PSD) at each electrode (within a 2 second moving window). This analysis confirmed that the human brain’s neural activity PSD is heterogenous, exhibiting a distinct topography with bilateral symmetry, consistent with prior resting-state MEG and iEEG studies ^26–28^.

However, investigating the variability of each region’s PSD over time, we identified that PSDs had the tendency to switch back and forth between two distinct spectral distributions. Moreover, these PSD switching patterns were found to form synchronous groups across multiple electrode locations, delineating large-scale functional networks, and in some cases connecting regions with distinctly different baseline PSDs.

Quantifying spontaneous naturalistic behaviors using simultaneous video and audio recordings of each subject, we found that the dynamic PSD switching of a subset of networks aligned with multiple observed spontaneous behavioral transitions. These network-behavior relationships were stable over multiple days, but were altered during sleep, suggesting state-dependent plasticity of brain-wide network organization.

Our results provide robust evidence for the presence of multiple synchronous neuronal networks across the human brain. The real-time PSD switching dynamics of these networks provide physiologically interpretable read-outs, demonstrating the parallel engagement of multiple brain regions in a range of concurrent naturalistic behaviors.

## Results

iEEG and audio-visual recordings were simultaneously obtained from pre-surgical, in-hospital, drug-resistant epilepsy patients (**Fig. 1**). For this study, we analyzed recordings obtained from 10 subjects implanted with bilateral depth electrodes that covered the frontal, temporal and partially occipital lobes (supplementary **Table S1**). The number of electrodes and the exact coverage varied among subjects to meet their clinical requirements.

**Fig. 1.**
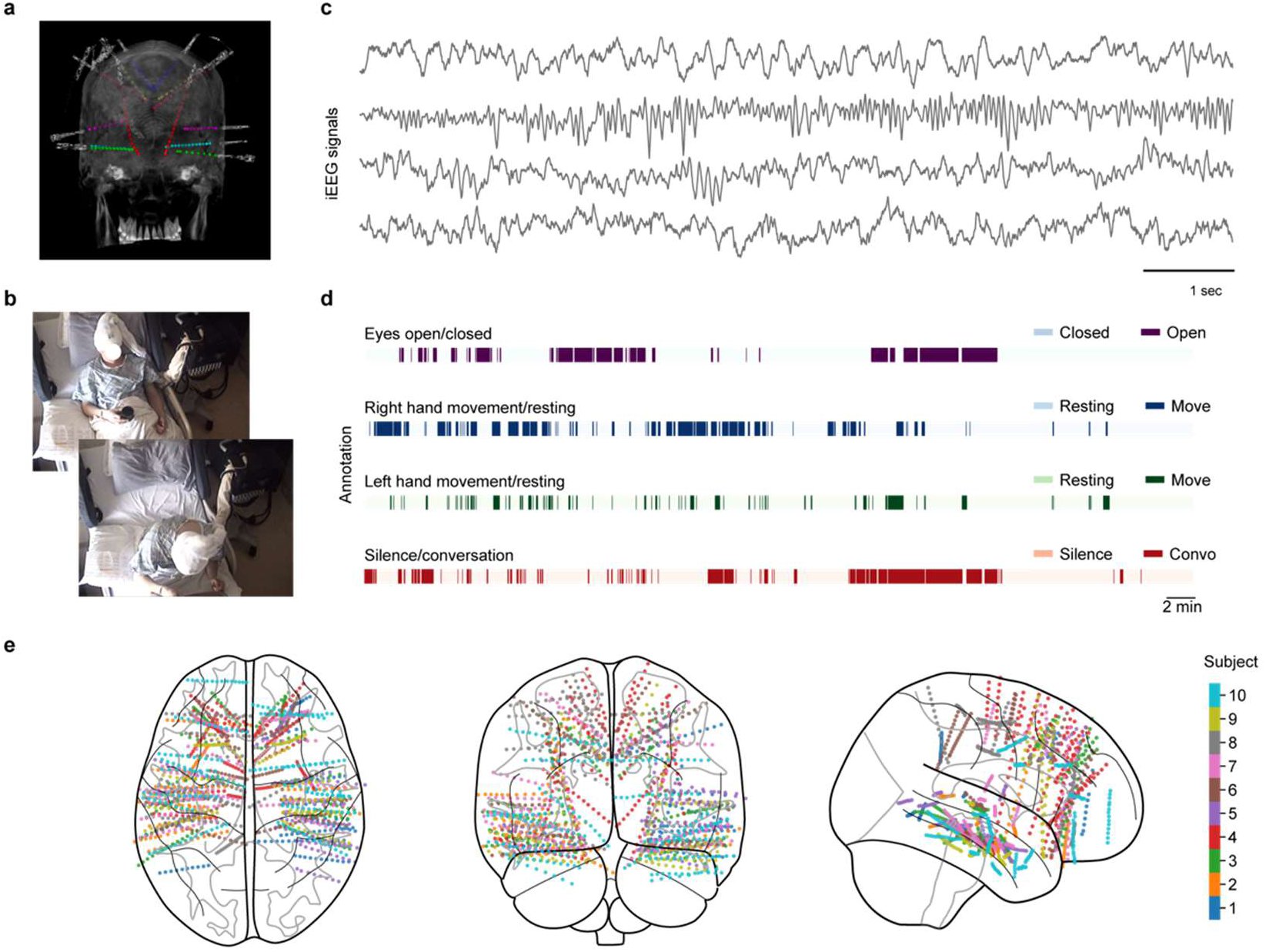
Experimental setup and overview of data sources. **a**, Illustration of the implant in an example subject with bilateral depth electrode arrays. **b**, Example frames from monitoring camera with patient leaning back and sitting forward. **c**, Example iEEG signals. **d**, Example annotation for different behavioral state transitions occurring simultaneously during one awake 60-minute recording. **e**, Illustration of the distribution of the implant in individual subjects on brain templates (axial, coronal, and sagittal views).

For each subject, electrode locations (MNI coordinates) were determined through the Intracranial Electrode VISualization (iELVis) toolbox^29^, which involved co-registering pre– and post-implant MRI and CT scans. The anatomical location of each electrode was then determined by mapping MNI coordinates onto the Automated Anatomical Labeling (AAL) brain atlas^30^ (see **Methods**). Real-time spontaneous naturalistic behaviors were evaluated by tracking the movement of the left and right hands using DeepLabCut^31^ (DLC) analysis, as well as by examining the spectral power of audio recordings. Behavioral state transitions were validated through manual video review of the recordings during the awake periods. Additional behavioral state transitions, such as eyes opening and closing, were also manually annotated for the same 60-minute wakefulness recordings (**Fig. 1d**).

### Time-averaged PSDs reveal spatially-varying distribution in the human brain

Moving-window PSD profiles for each electrode were calculated using 2-second epochs with 50% overlap. The 1/f component of broadband PSDs (2-55 Hz) was estimated and removed using the FOOOF algorithm^32^ within each sliding window. We focused on the 2-55 Hz frequency range to minimize noise and edge effects.

For each subject, we chose a period of 5 minutes of continuous ‘resting state’ data (in which the subject sat quietly making no discernable movements). The time-averaged PSD over this period was calculated for each electrode channel and aggregated across subjects by mapping the anatomical location of every electrode using the AAL brain atlas. **Fig. 2a and b** (i) depict the resulting PSD distributions, which are heterogeneous across the brain, exhibiting bilaterally symmetric organization with natural regional subdivisions (supplementary **Fig. S1a**). Peak powers at lower frequencies are seen in the mesial and lateral temporal regions, while peak powers at higher frequencies are seen in the frontal and anterior superior parietal regions (**Fig. 2b**).

**Fig. 2.**
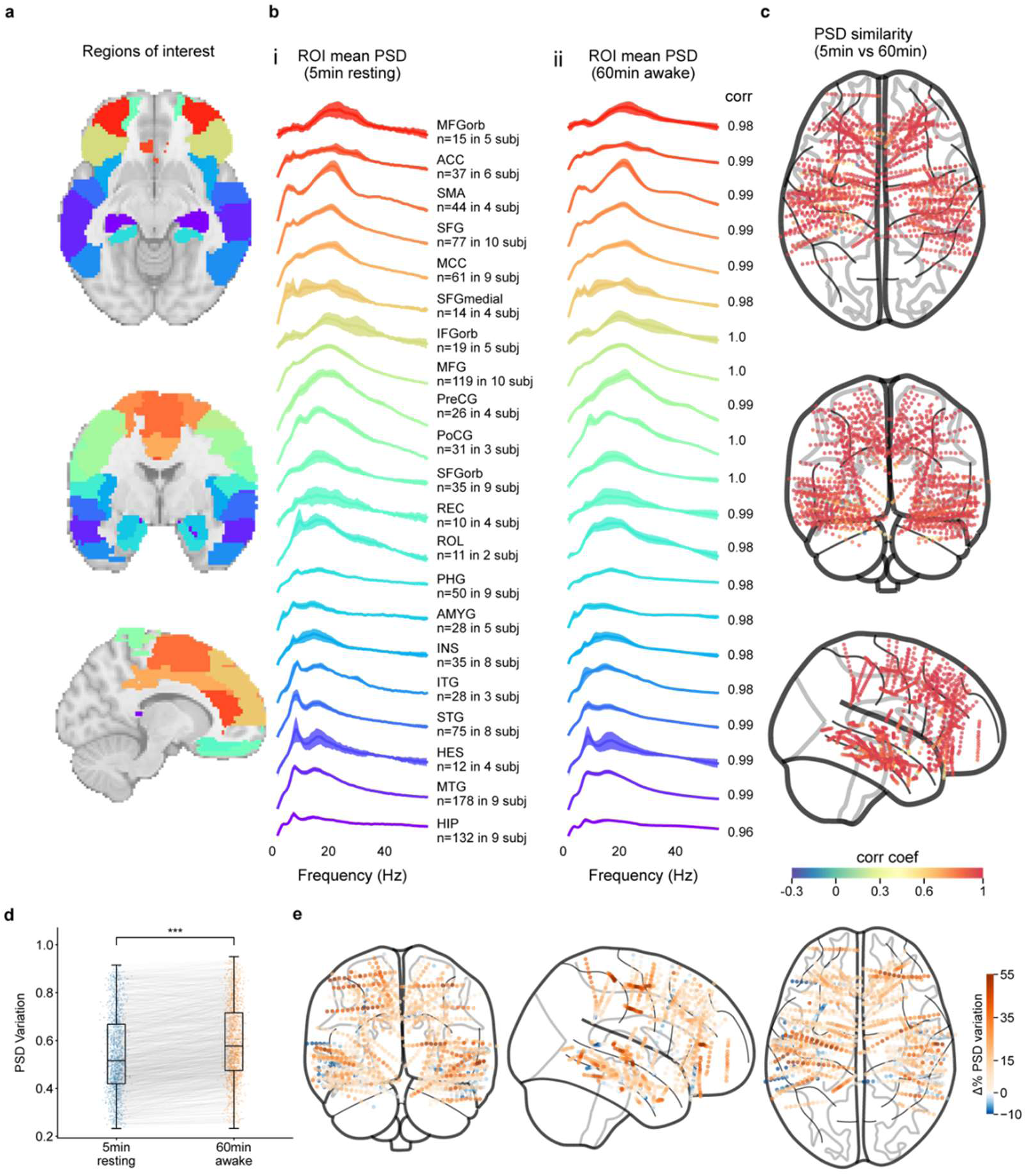
Spatial spectral architecture in the human brain. **a**, Pre-selected regions of interest demarcated by color coding, as shown in subplot **b**. **b**, PSD profiles averaged across all recording sites within each ROI (mean±2.58 SEM) during the 5-min resting period (i) and during the 60-min wakefulness period (ii), ranked and color-coded by peak frequency. **c**, Similarity of time-averaged PSD profiles between the 5-min resting and 60-min wakefulness in individual channels on a brain plot, where color indicates Pearson correlation coefficient (r=0.928±0.14, mean±SD, n=1513). **d**, Distribution of PSD variation during the 5-minute resting period vs. 60-minute wakefulness. Significantly greater variation was found during 60-min wakefulness (p ≤1×10^−6^, n=1513, Wilcoxon signed-rank test, one-sided). **e**, Percentage of change in PSD variation between 5-min resting and 60-min wakefulness in each individual channel on a brain schematic. In the box plot **d**, the box extends from the first quartile to the third quartile of the data, with a line at the median. The whiskers extend from the box to the extreme data points. All data points including outliers are shown individually.

**Fig. 2b** (i-ii) compares these resting time-averaged PSD distributions to PSDs averaged over 60 minutes of continuous data in which the subject was actively moving and interacting with their environment. Strong similarity is seen between resting and awake states (**Fig. 2c**).

We next compared the variability of moving-windowed PSDs over time for the same awake and resting epochs (supplementary **Fig. S1b**). The 60-minute awake period demonstrated significantly more variability compared to the 5-minute resting state (**Fig. 2d-e**). These results suggest a relatively stable architecture of PSD profiles across the human brain that reflects intrinsic characteristics of activity at each cortical location across both resting and awake states. However, increased variability in PSDs during wakefulness suggests that this overall pattern is not static.

### PSDs exhibit switching behavior between 2 states

**Fig. 3a** shows an example moving-window PSD sequence (2-second length, 50% overlap) for an electrode channel located in the left superior temporal gyrus of Subject 6. The spectral shape of the PSD can be seen to switch back and forth over time. Applying K-means clustering to these PSDs, we found two distinguishable PSD clusters for this channel, as plotted in **Fig. 3b** (i).

**Fig. 3.**
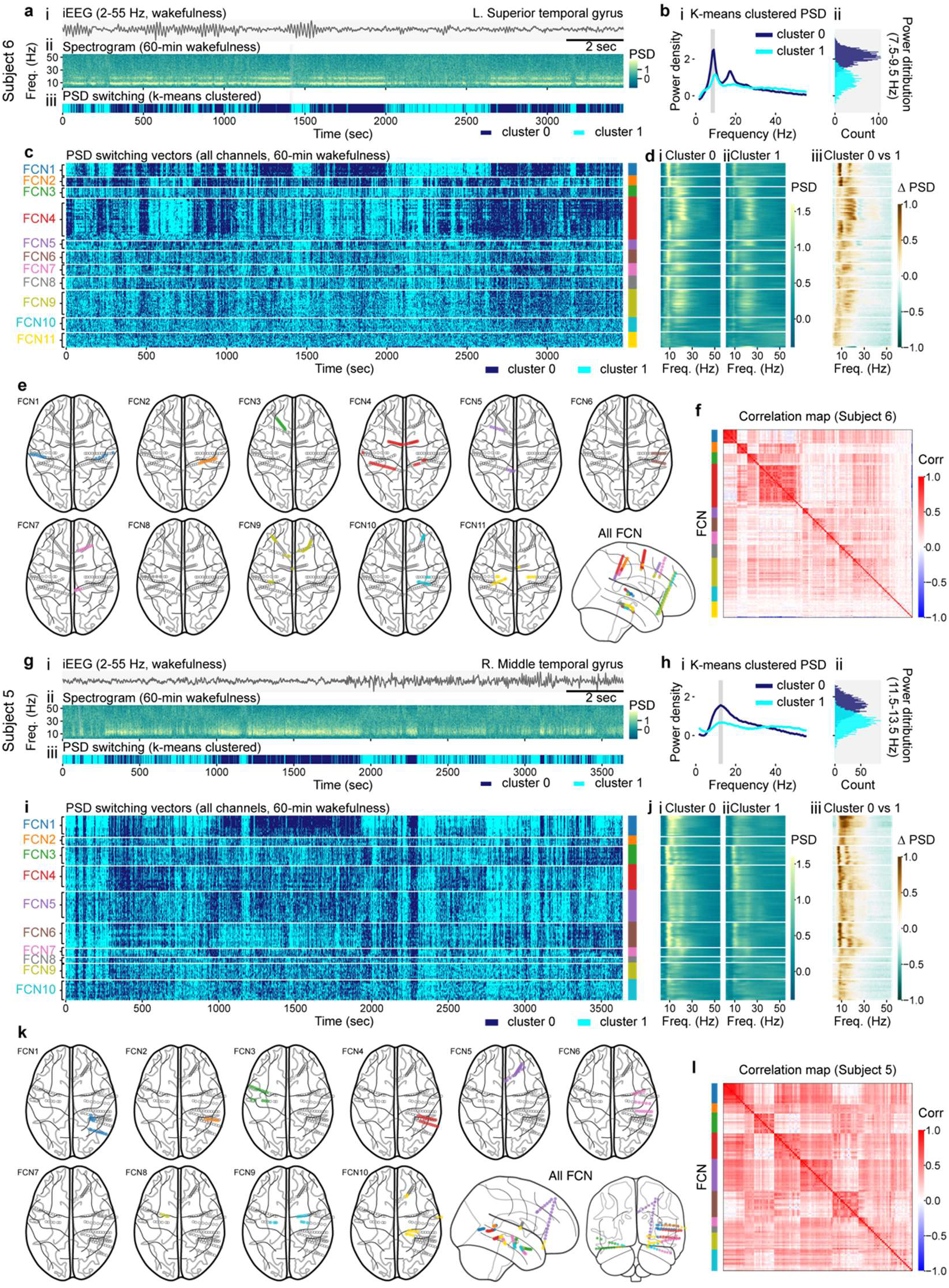
PSD profile clustering and switching demonstrates evidence of large-scale functional. **a**, (i): iEEG recordings of an example channel located in the superior temporal gyrus from Subject 6. (ii): Whitened sliding-windowed PSDs from the same channel during the 60-min wakefulness. (iii): K-means clustering of sliding-windowed PSD profiles shown above. **b**, (i): Mean PSD profile of each cluster (mean±1.96 SEM). (ii): Distribution of the power averaged between 7.5-9.5 Hz (p≤1 × 10^-6^, Mann-Whitney U test, two-sided), color-coded by the K-means clustering results. **c**, PSD switching vectors of each individual channel in Subject 6 (blindly from behavioral information), sorted by the network clustered through hierarchical clustering. **d**, Mean PSD profiles of clusters 0 (i) and 1(ii) in individual channels, with channels listed in the same order as shown in subplot **c**. (iii): Differences in PSD profiles between the two clusters. **e**, Location of channels in each network of Subject 6. **f**, Correlation map of PSD switching vectors in Subject 6. **g-l**, Results from Subject 5, following the same configuration as shown in subplot **a-f**.

The time-varying switching between these two PSDs is represented in binary form in **Fig. 3a** (iii) where the cluster with smaller gamma band power (35-55 Hz) was arbitrarily defined as cluster 0. This approach was extended to all channels in Subject 6 by independently computing 2 PSD clusters for each electrode. **Fig. 3c** displays the binary PSD switching vectors for all channels, along with the mean PSD profiles in each cluster (**Fig. 3d**). Examining the change in PSD for each channel (**Fig. 3d** (iii)), we see that electrode-specific switching generally corresponds to reciprocal decreases/increases in power at low frequency bands (e.g. theta (4-8 Hz), alpha (8-12 Hz) or beta (12-30 Hz) bands) v/s higher power bands (e.g. low gamma (30-55 Hz)). We also note that for a range of intermediate (channel-specific) frequencies, no apparent change in power would be observed during switching. Equivalent analysis is applied to data from a second subject in **Fig. 3g-j**. In both subjects, a range of different switching patterns can be seen to occur across all of the channels, with certain subsets exhibiting temporally coherent switching behavior.

The presence of 2 PSDs for the epoch in **Fig. 3a** is confirmed in **Fig. 3b** (ii), which finds 2 distinct distributions of PSD spectral power at 7.5-9.5 Hz (the frequency that explains the most variance in this channel). Across subjects and channels, the presence of at least 2 clusters was confirmed using the Duda-Hart test (whether a data set should be split into 2 clusters) (**Fig. S2 a**).

### PSD switching synchrony reveals multiple functional networks

The binary representations of PSD switching across all electrode channels in **Fig. 3c and i** were sorted into correlated groups using hierarchical clustering, with the optimal number of clusters for each subject determined using silhouette score (Supplementary **Fig. S2 c**). The resulting groups were sorted vertically based on their high-to-low mean within-cluster correlation (see **Methods**). In both examples, the top groups of channels exhibit distinct PSD switching patterns over time.

**Fig. 3e and k** projects all channels onto their anatomical positions in the brain, showing that grouped channels represent correlated networks that span multiple brain regions including bilateral homologous areas. Correlation matrices comparing PSD switching vectors across channels are shown in **Fig. 3f and l**, demonstrating additional interrelationships between each network group. These results demonstrate that multiple electrodes across the brain exhibit highly correlated PSD switching behaviors, consistent with inter-regional communication and functional connectivity.

Importantly, we note that these groupings, derived based on correlated PSD switching dynamics, include channels that have distinctly different baseline PSD profiles. For example, in subject 6 functional networks (FCNs) 4 and 5 both contain at least two subgroups of channels with distinctly different PSD profiles, but correlated PSD switching vectors (**Fig. 3c**). Further examination of these subsets confirmed that FCN4 channels within the supplementary motor area correspond to the subgroup with a PSD peak in the beta band, while channels located in the postcentral gyrus correspond to the subgroup with a PSD peak in the alpha band. Similarly, in FCN 5, channels whose PSDs peak in the alpha band are located in the posterior cingulate gyrus, whereas PSDs peaking in the beta band are located in the middle frontal gyrus. Additional examples of coherent PSD switching between brain regions with differing baseline PSDs can be observed in FCN 6 and 9 in subject 6, and FCN 1,4, and 6 in subject 5 (**Fig. 3d and j**, and **Fig. S2d**).

This analysis approach effectively captures variations in neural activity across a broad range of frequencies, while reducing the high-dimensionality of neural activity to a dynamic binary representation of a PSD switching vector. In turn, the synchronization of these PSD switching dynamics between diverse brain regions reveals multiple long-range functional networks. Applying this network discovery approach to all subjects, we found 2 to 11 networks per subject (dependent on patient-specific electrode coverage) as summarized in supplementary **Fig. S3**. Across subjects, networks were found to span both short and long-range brain regions, and include areas with varying baseline PSD profiles, providing a novel view of dynamic coordination between distinctly different brain areas.

### Multi-threaded network PSD switching correlates with behavioral dynamics

All PSD switching was characterized from real-time neural recordings, independently of behavioral information. However, for each subject we also quantified a range of real-time naturalistic behaviors (eyes open/closed, conversation/silence, left or right-hand movement/resting) with manual annotation (see **Methods**).

Comparing these behavioral state transitions to PSD switching dynamics for the same recording period, we noticed striking agreement between specific behaviors and the dynamics of switching in individual functional networks. For example, **Fig. 4a** (i-iii) shows quantified spontaneous behaviors (right and left-hand movement/resting, conversation/silence) for subject 6, alongside the PSD switching vectors of channels within the clustered networks that had the highest Pearson correlation to each behavior. Equivalent results for a second subject are shown in **Fig. 4f** (i-iii), for eyes open/closed, left-hand movement/resting and conversation/silence.

**Fig. 4.**
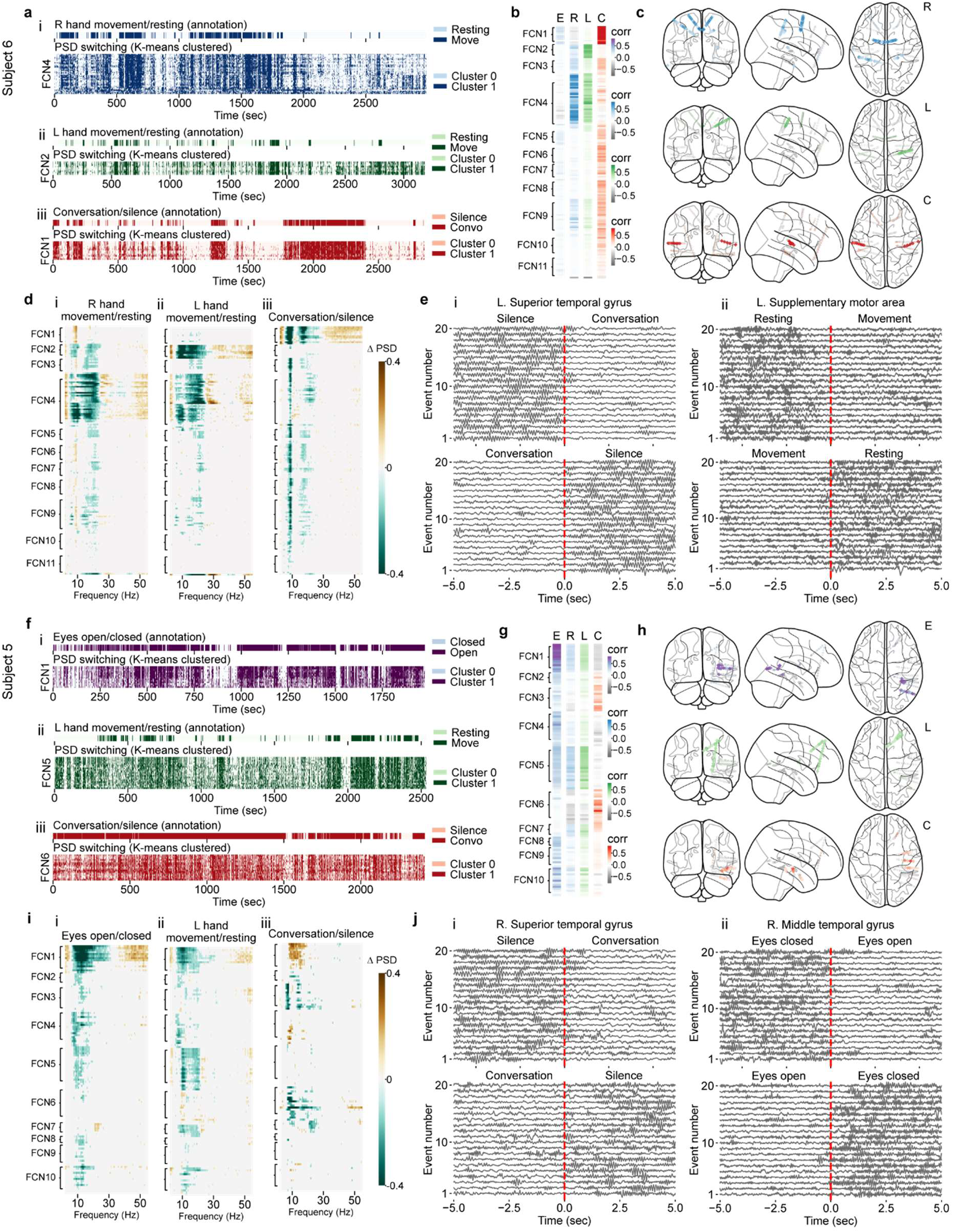
PSD switching correlates with behavioral state transitions. **a**, Annotated behavioral state transitions and corresponding PSD switching vectors from responsive networks for right-hand movement – resting (i), left hand movement-resting (ii) and conversation-silence (iii). **b**, Correlation coefficients of individual channels in response to each category of state transition (E: eyes open –closed, R: right-hand movement-resting, L: left-hand movement-resting, C: conversation-silence), sorted by networks. **c**, Spatial distribution of correlation coefficient on brain templates, with colors indicating the Pearson correlation to the presented behavioral category. **d**, PSD profile differences in individual channels across state transitions of right-hand movement-resting (i), left hand movement-resting (ii) and conversation-silence (iii), listed by networks. Color indicates statistically significant changes (p≤0.05, Mann-Whitney U test, two-sided, FWER corrected). Dominant statistically significant changes for each behavior fall within our previously-defined networks (FCN1, 2 and 4). **e**, (i): iEEG recordings in an example channel located at the left superior temporal gyrus during 20 events from silence to conversation and 20 events from conversation to silence. (ii): iEEG recordings in an example channel located at the left supplementary motor area during 20 events from resting to right-hand movement and 20 events from right-hand movement to resting. **f-j**, Examples from subject 5, following the same configuration as shown in subplot **a-e** with behavioral state transition across eyes open/closed, left-hand movement/resting and conversation/silence.

Strong agreement is seen between the dynamics of specific networks and categories of behavioral state transitions. Moreover, it is important to note that all variables are plotted in real-time, simultaneously for the same 60-minute epoch. This means that the behaviorally-linked PSD-switching dynamics within each of these networks are all occurring at the same time, representing multiple threads of network activity operating in parallel and associated with different facets of ongoing naturalistic behavior.

Correlations between observed behavioral state dynamics and the PSD switching vectors for each individual electrode channel are shown in **Fig. 4b and g** for subjects 6 and 5 respectively (with channels ordered by their networks as in **Fig. 3c and i**). **Fig. 4c and h** projects these channel-specific correlation values onto the spatial location of every electrode. The behavior-correlated dynamic networks revealed again span brain-wide and bilaterally symmetric regions (where bilateral electrodes are present).

It is important to note that the spatial localization of behaviorally-correlated channels spans both primary and associative areas corresponding to each respective behavioral category. For example, networks responsive to alternating conversation/silence epochs (FCN 1 in subject 6 and FCN6 in subject 5) include channels located in Heschl’s gyrus, middle and superior temporal gyrus, recognized as primary and associative auditory and language areas. Networks responsive to left-hand movement/resting (FCN2 in subject 6 and FCN5 in subject 5) include channels located in the postcentral gyrus, middle and superior frontal gyrus, and middle cingulum, while the network responsive to right-hand movement/resting (FCN4 in subject 6) includes channels located in same regions but also supplementary motor area. These areas correspond to contralateral primary and supplementary motor, primary somatosensory cortex, as well as areas associated with motor planning, execution and coordination. The network responsive to eyes open/closed (FCN1 in subject 5) includes channels located in right middle temporal gyrus (posterior), a site associated with visual information processing^33^.

The spectral ranges for each channel that exhibit significant decreases or increases between ‘active’ v/s ‘inactive’ states are shown for each behavior type in **Fig. 4d and i**. Independent of our PSD switching analysis, these results identify channels with clear relationships between behavioral state changes and diverse changes in spectral power. Importantly, channels with similar behavioral tuning fall within our previously delineated functional networks. We also note that the differing frequency-dependence of increases, decreases or minimal changes observed within each channel would make network synchrony almost impossible to discern in observations that were limited to a narrow frequency range.

Examples of broadband iEEG samples (2-35 Hz) from channels within correlated networks for epochs spanning 5-seconds before and after the specified behavioral state transition are shown in **Fig. 4e and j** (20 events from inactive to active and 20 events from active to inactive). Rapid changes in iEEG signals are observed at the onset and offset of each behavior, consistent across individual events, indicating the temporal sensitivity of iEEG signals to behavioral state transition.

### Mapping of brain-wide network activity to behavioral states across subjects

**Fig. 5** consolidates data from all subjects, mapping the correlation coefficient between each behavior and each channel’s PSD switching vector (**Fig. 5a** (ii)) onto their corresponding brain location (**Fig. 5b**). Significant changes in each channel’s PSD between cluster 1 vs 0 is shown in **Fig. 5a** (i). Channels from all subjects are ordered by their anatomical brain region.

**Fig. 5.**
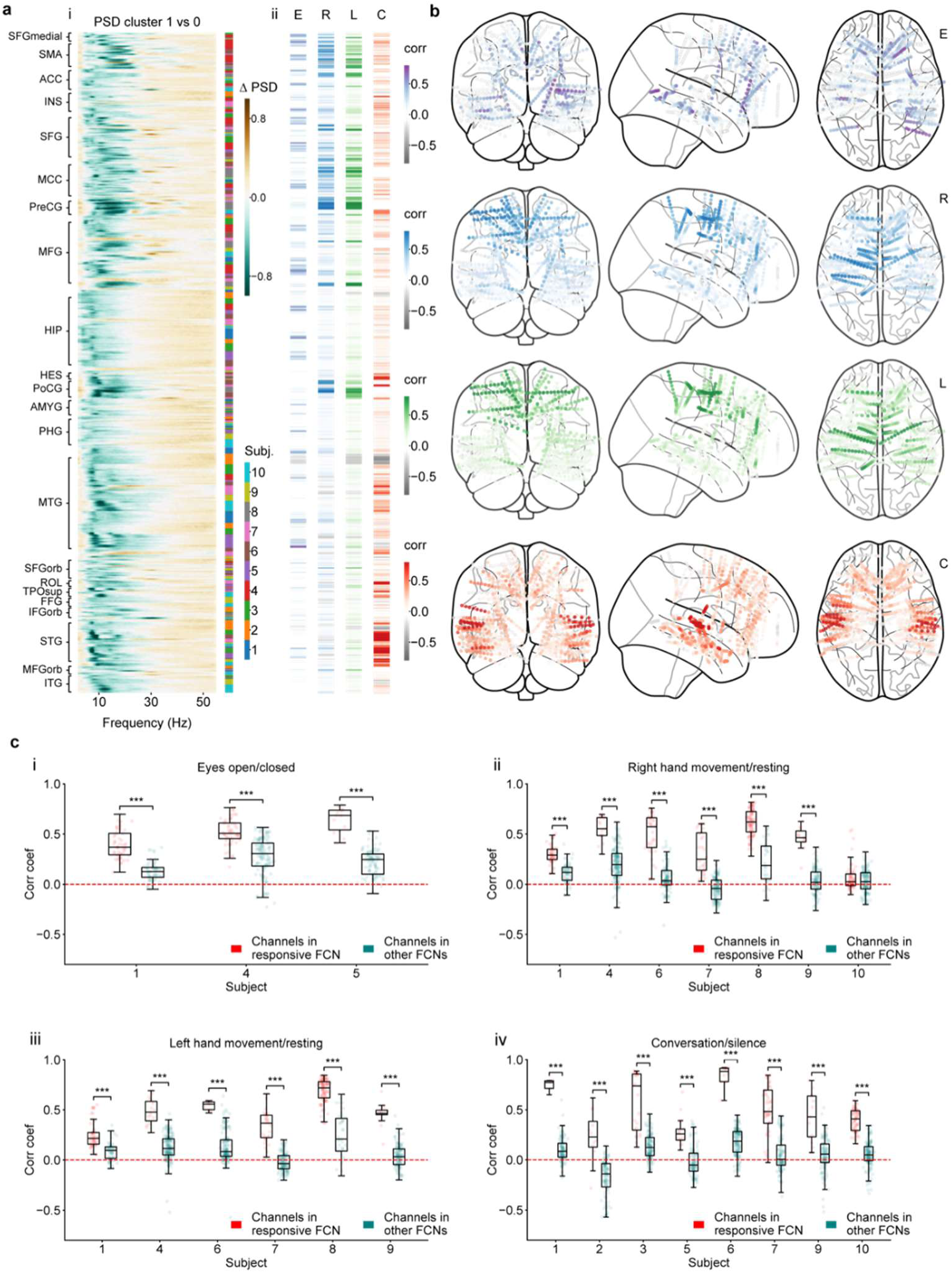
Responsive channels and networks across subjects. **a**, (i): Differences between PSD profiles averaged within cluster 0 and 1 in individual channels across all subjects, sorted by brain areas. (ii): Correlation coefficients of listed channels in response to each category of behavioral state transitions. **b**, Spatial distribution of correlation coefficient of individual channels on brain template across subjects in response to eyes open/closed (N=7), right-hand movement/resting (N=10), left-hand movement/ (N=10) and conversation/silence (N=9), from top to bottom. Channels from subjects with not enough length of valid annotation are marked as hollow dots. **c**, Distribution of correlation coefficients in channels from the responsive networks and other networks within each subject. (i): eyes open/closed (p≤1 x10^−6^ in 3 subjects). (ii): right-hand movement/resting (p≤1×10^−6^ in 6 subjects, p>0.05 in 1 subject). (iii): left-hand movement/resting (p≤1×10^−6^ in 6 subjects). (iv): conversation/silence (p≤1×10^−6^ in 8 subjects). All tests were conducted using the Mann-Whitney U test, two-sided, with FWER corrected. In the box plots **c** (i-iv), the box extends from the first quartile to the third quartile of the data, with a line at the median. The whiskers extend from the box to the extreme data points. All data points including outliers are shown individually. Asterisks show significance level of *p≤0.05, **p≤0.01 and ***p≤0.001.

A general pattern of power suppression in the low frequency range (4-30 Hz) and an increase in gamma band power (30-55 Hz) was observed across individual responsive channels (r>=0.5), although with regional variations. As summarized in **Table S4**, across all subjects, observed responsive regions again span multiple lobes and both hemispheres, including brain regions involved in both primary and secondary processing.

In **Fig. 5c** (i-iv) we confirm the selectivity of PSD switching dynamics to specific behavioral categories by comparing the correlation of channels from the most tuned network to the correlation of channels in all other networks to each behavioral category, for each subject. In all cases, the correlation coefficients within the most responsive network were significantly greater than correlation coefficients for the other networks.

### Comparing functional networks across days and wake / sleep states

The results above used a range of different data-driven approaches to identify brain networks whose broadband PSD switching dynamics exhibited surprising correlations to observable transitions in behavioral state. To cross-validate our results and explore the stability of this network organization over time, we repeated analysis on datasets acquired across multiple days, as well as comparing awake periods to recordings during N3 sleep state for each subject.

The N3 sleep state was extracted from the full night sleep data of each subject based on the beta/delta power ratio (see **Methods** and supplementary **Fig. S5**). To track behaviors across subjects, we implemented a semi-automated approach; first extracting the audio spectrogram from surveillance videos, and second, tracking the movement of the left and right hands using DeepLabCut^31^. Changes in silence/conversation states were approximated using audio power within 80-400 Hz range, covering the human vocal frequency range. Left/right hand movement are approximated by computing the motion energy based on DeepLabCut body parts labeling. This approach allowed us to analyze large scale datasets while reducing potential bias due to subjective errors during video annotation.

**Fig. 6** compares PSD switching dynamics for Subject 6 for two awake periods (on consecutive days) and a period of N3 sleep (**Fig. 6c,d,e**). All channels are ordered vertically into networks using hierarchical clustering of the PSD switching binary vectors from awake day 1. The presence of similarly switching groups of channels within each network on days 1 and 2 are clearly visible, confirming the stability of basic network organization between days. The two PSD clusters identified for each channel (**Fig. 6c-d** (v)) were also remarkably similar between day 1 and 2 awake datasets. The correlation matrix of PSD switching vectors between days 1 and 2 are also consistent. The specialization of each channel / network to each of the three quantified behavioral states (right– and left-hand movement, and conversation) is also maintained. As expected, there was no correlation between the real-time PSD switching vectors between days 1 and 2, nor between real-time behavioral state dynamics.

**Fig. 6.**
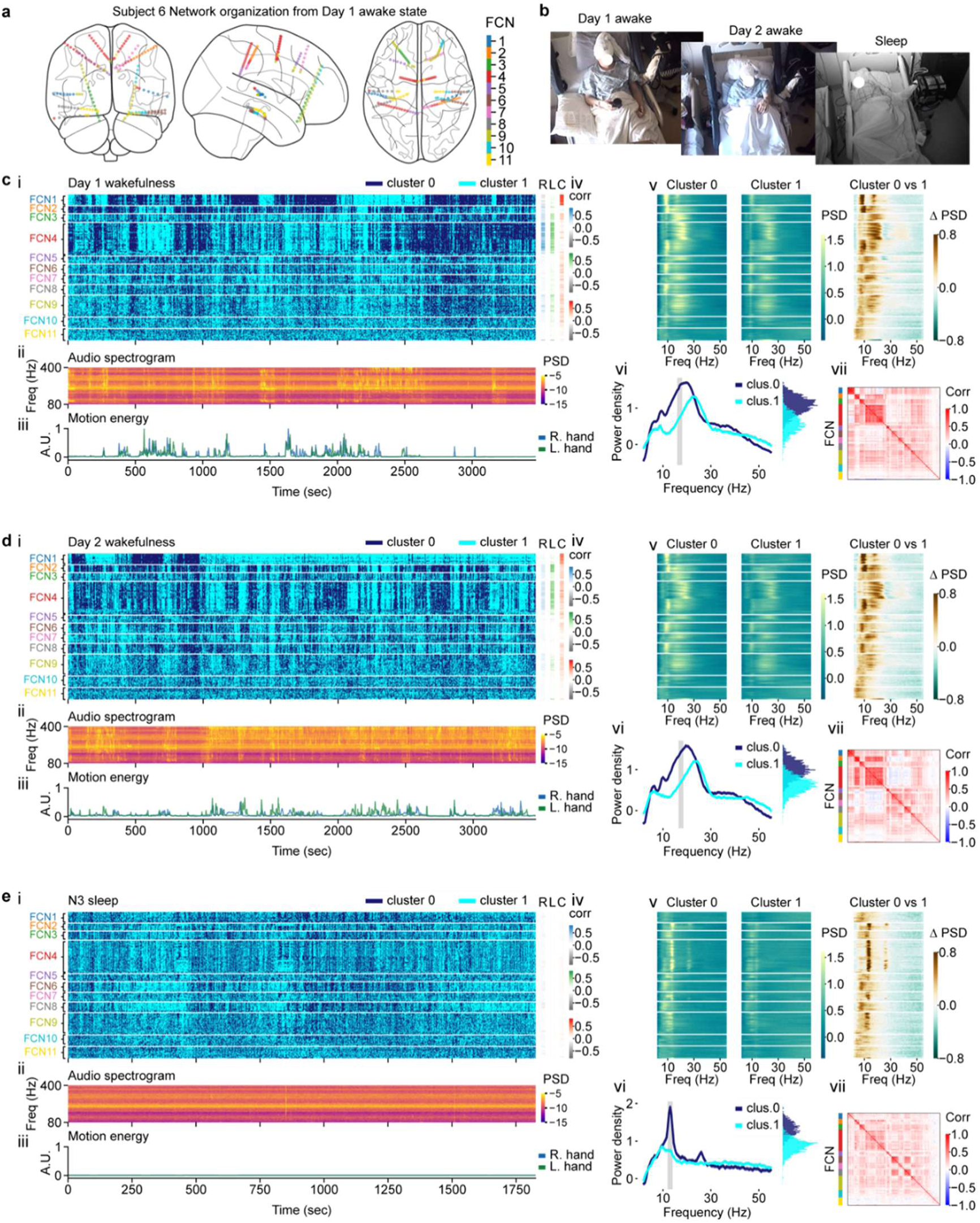
Comparing network dynamic results across days and sleep state (subject 6). **a**, Network organization characterized from day 1 wakefulness. **b**, Screenshots of monitoring video from day 1 awake, day 2 awake and N3 sleep period. **c**, (i): PSD switching vectors of individual channels during day 1 wakefulness, listed by networks. (ii): Audio spectrogram from simultaneous audio-visual recordings. (iii): Motion energy of DLC-labeled left/right-hand positions from simultaneous audio-visual recordings. (iv): Correlation coefficients of individual channels in response to each category of behavioral changes (right (R) and left (L) hand motion energy, and audio power (C)). (v): Mean PSD profiles of clusters 0 (left) and 1 (middle) from K-means clustering in individual channels, and their differences (right), listed by networks shown in subplot **c** (i). (vi): Mean PSD profiles (mean±1.96 SEM) of each cluster from an example channel located in left supplementary motor area, along with distribution of power averaged between 16-18 Hz (p≤1×10^−6^ Mann-Whitney U test, two-sided), color-coded by the K-means clusters. (vii): Correlation matrix of PSD switching vectors of individual channels. **d**, Results from day 2 wakefulness, following the same configuration as shown in subplot **c**. **e**, Results from N3 sleep state, following the same configuration as shown in subplot **c**. **c**(i) intra-network correlation: r=0.3±0.17, mean±SD, n=11 networks, (Pearson correlation), change in inter-network correlation: p≤1×10^−6^, n=165, Wilcoxon signed-rank test, one-sided, adjusted inter-network distance (see **Methods**). Pearson correlations between awake day 1 and 2 PSDs in **c-d(v)**: for PSD 0: r=0.98±0.02; and PSD 1: r=0.94±0.07, between awake day 1 and N3 sleep: PSD 0: r=0.65±0.18; and PSD 1: r=0.26+0.31, mean±SD, n=165 channels. Pearson correlation between awake day 1-2 PSD vector correlation matrices **c-d(vii)**: r = 0.75, between day 1 to N3 sleep (**c-e(vii)**) r = 0.44.

The same analysis repeated on data acquired in the same subject during N3 sleep, reveals markedly different patterns (**Fig. 6e**). First, we see that the two PSD profiles found by initial clustering of moving window PSD data are distinctly different from those found in awake datasets. Noting that consistent vertical ordering (and thus network grouping) was used for days 1 and 2, and the N3 sleep data, we observe decreased similarity between PSD switching vectors within network groups (intra-network correlation). However, we also see increased similarity across network groups (inter-network correlation).

Importantly, despite the lack of behavioral state dynamics (movement or conversation), distinct patterns of PSD state switching are still visible during N3 sleep, and span across networks that had uniquely different dynamics in the awake state. The correlation matrix of PSD switching vectors from N3 sleep (**Fig. 6e** (vii)) confirms this more uniformly correlated pattern overall, while being distinctly different relative to day 1. Additional examples from other subjects are shown in supplementary **Fig. S6** and **Fig. S7.**

### Consistency and plasticity in network organization

**Fig. 7** expands the analysis above to all subjects, examining and comparing awake datasets acquired on two consecutive days, as well as data acquired during N3 sleep. **Fig. 7a** shows that across subjects, regional baseline PSD profiles agree on consecutive days for awake periods, while PSD profiles during N3 sleep demonstrated significantly decreased correlation compared to day 2 awake.

**Fig. 7.**
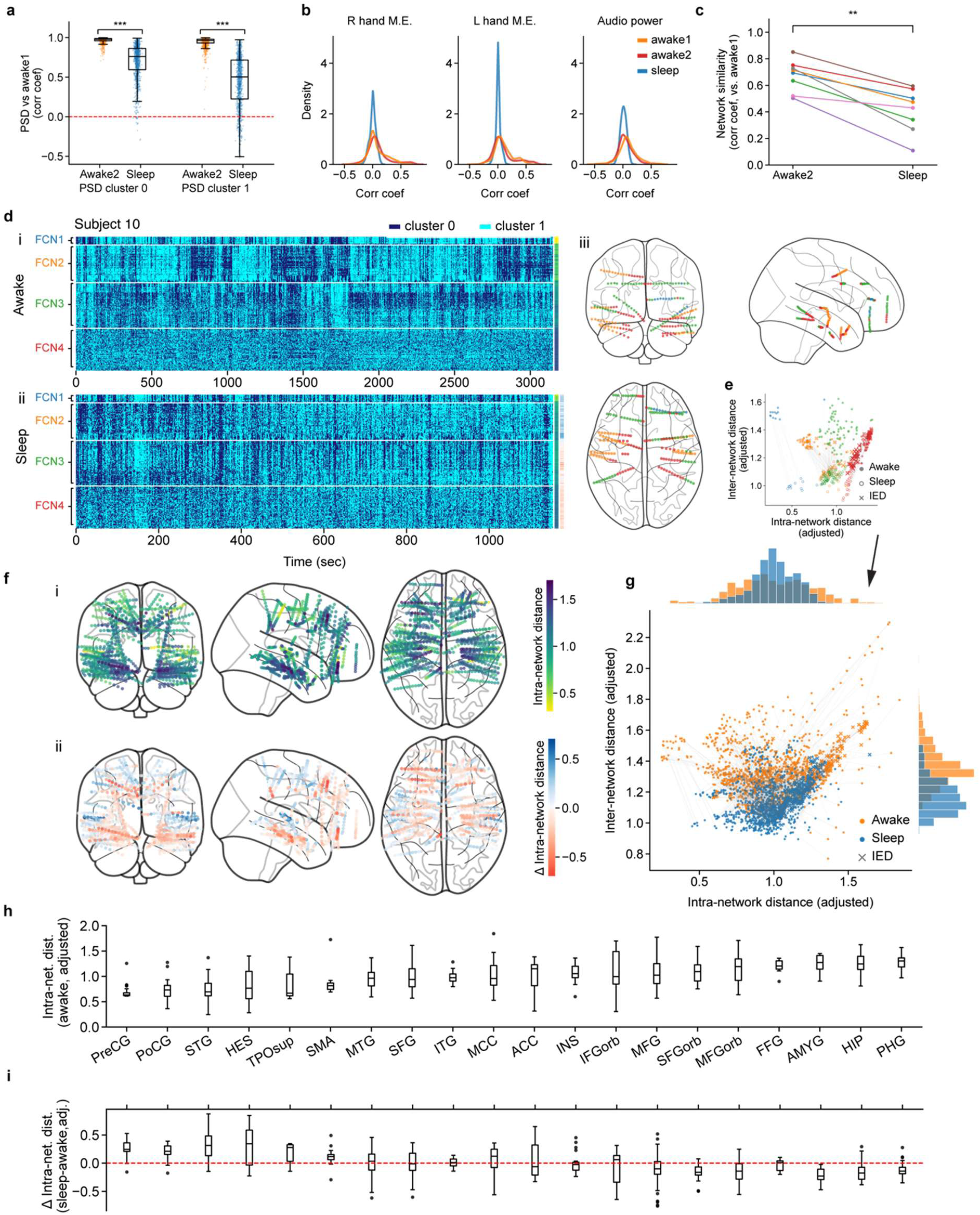
Network organization comparison across different recordings. **a**, Comparison of PSD profiles’ similarity to those characterized day 1 wakefulness (day 2 awake cluster 0: r= 0.97±0.03, cluster 1: r= 0.94±0.07; sleep cluster 0: r=0.69±0.22, cluster 1: r=0.44±0.33. n=1170, mean±SD, Pearson correlation). PSD profiles from N3 sleep show significantly less similarity than those from day 2 wakefulness (cluster 0: p≤1 × 10^-6^, cluster 1: p≤1 × 10^-6^, n=1170, Mann-Whitney U test, one-sided). **b**, Correlation between PSD switching vectors to extracted right-, left-hand motion energy and audio power. PSD switching vectors characterized from N3 sleep demonstrate ∼zero correlation with behavioral changes (r=0.002±0.06, r=0.006±0.04, r=-0.009±0.05, n=1170 channels, mean±SD, Spearman Rho). **c**, Connectivity matrix similarity compared to those characterized day 1 wakefulness (day 2 awake: r=0.67±0.11; N3 sleep: r=0.41±0.15, n=8, mean±SD, Pearson Correlation). Connectivity matrices observed from N3 sleep demonstrate significantly less correlation than day 2 wakefulness (p=0.004, n=8, Wilcoxon signed-rank test, one-sided). **d**, PSD switching vectors of individual channels from subject 10 during day 1 awake (left) and N3 sleep state (right), sorted by networks characterized from day 1 wakefulness. **e**, Adjusted intra-network distance versus inter-network distance for each individual channels during day 1 wakefulness and N3 sleep in subject 10, color indicates the network groupings as shown in **d**. **f**, Spatial distribution of adjusted intra-network distance during day 1 wakefulness (i), and variations in adjusted intra-network distance between N3 sleep and day 1 wakefulness (ii). **g**, Adjusted intra-network distance versus inter-network distance for each individual channels during day 1 wakefulness and N3 sleep across all subjects (N=8). Significant decrease in the mean of adjusted inter-network distance (p≤1×10^−6^, n=1176, Wilcoxon signed-rank test, one-sided) and decrease in the variance of adjusted intra-network distance (p≤1×10^−6^, n=1176, F-test, one-sided) were observed during sleep state. **h**, Distribution of adjusted intra-network distance within each brain area during day 1 wakefulness, ranked from the lowest to the greatest. **i**, Distribution of change in adjusted intra-network distance within each brain area between day 1 wakefulness and N3 sleep state, displayed in the same order as shown in **h**. In the box plot **a**, **h** and **i**, the box extends from the first quartile to the third quartile of the data, with a line at the median. The whiskers extend from the box to the extreme data points. All data points including outlies are shown individually in **a**. Outliers are shown as fliers in the box plots **h** and **i**. Asterisks show significance level of *p≤0.05, **p≤0.01 and ***p≤0.001.

The distributions of correlation coefficients between PSD switching vectors and each behavior across all channels are similar for awake days 1 and 2, but demonstrate ∼zero correlation during N3 sleep (**Fig. 7b**). Supplementary **Fig. S8** maps correlations between PSD switching and awake behaviors for individual channels across subjects, demonstrating relatively stable representations between days 1 and 2. **Fig. 7c** demonstrates the strong similarity between correlation matrices of PSD switching vectors across channels for awake day 1 and 2, as well as the significant dissimilarity between awake day 1 and N3 sleep. This reduced network similarity suggests plasticity of the underlying network structure.

A further example of PSD switching vector network clusters is shown in **Fig. 7d** comparing day 1 awake and N3 sleep states for Subject 10. We again see that in the awake state, channels in networks 1-3 show PSD switching that is synchronous with other channels within their network (intra-network), but asynchronous with channels in other networks (inter-network). During N3 sleep, the same channels reduce their intra-network synchrony, while increasing their inter-network synchrony. Importantly, we also note that network 4, which has low intra-network synchrony in the awake state, exhibits a significant increase in synchrony within its network, and the rest of the channels during N3 sleep.

To quantify these effects, we computed the averaged inter-network and intra-network correlation distance for each channel in day 1 awake and N3 sleep states (normalized by the grand mean of respective intra-network distances to enable comparisons, see **Methods**). **Fig. 7e** and **Fig. 7g** plot how these parameters change from awake to sleep stated for all channels, in Subject 10 and across subjects respectively. Consistent decreases in inter-network distances (increased similarity across the brain) are seen during N3 sleep. We also see two distinct trends in intra-network distance; channels with low intra-network distance in the awake state tend to increase during N3 sleep, while channels with high intra-network distance tend to decrease towards a common middle state during N3 sleep, such that the overall variance of intra-network distances is reduced.

Focusing on channels with high intra-network distances in the awake state, we note that these channels are grouped within higher-numbered FCNs, and their PSD switching vectors appear more scrambled and random, which we refer to as “fast-switching”. Across subjects, these networks all exhibit a decrease in intra-network distance (increased brain-wide synchrony) during N3 sleep compared to the awake state (visible in **Fig. 7d-e**, **Fig. 6c-e, Fig. S6** and **Fig. S7)**.

Projecting channel-specific awake-state intra-network distances onto the brain for all subjects in **Fig. 7f** (i), we can see localization of channels with high intra-network distances (dark blue points) to the mesial temporal and frontal region, spanning parahippocampal gyrus, hippocampus, amygdala, middle frontal gyrus, orbital part in superior frontal and middle frontal gyrus, and fusiform gyrus (**Fig. 7h**). Projecting the net change in intra-network distance for all subjects between awake and N3 sleep states onto the brain in **Fig. 7f** (ii), we see that these same brain regions (**Fig. 7i**) exhibit the largest decreases in intra-network distance (increased synchrony) during N3 sleep (red points).

## Discussion

### Brain-wide PSD-switching dynamics reveal functional networks

This study utilized real-time brain-wide iEEG recordings in human subjects. Data was analyzed to seek representations of brain-wide neuronal networks, and to explore the relationship between the dynamics of network activity and real-time behavior. After first characterizing the distribution of PSD profiles across the brain, which generalized across subjects (**Fig. 2**), we recognized that PSDs at all recording sites had the tendency to switch back and forth between 2 apparent PSD states (**Fig. 3a,b/g,h**). Quantifying these real-time PSD switching dynamics for all electrodes, we discovered synchronies in switching, not just between adjacent channels, but also in distant or contralateral electrode locations (**Fig. 3c,e/i,k**). Investigating these groupings further, we found that synchronous switching could occur between channels whose underlying PSD spectral shapes were dissimilar, emphasizing that PSD switching can be coordinated between functionally different brain regions (**Fig. 3d,j)**. We propose that these switching synchronies constitute a representation of brain-wide functional networks.

### Distinct network dynamics correlate with concurrent naturalistic behaviors

PSD switching patterns within each putative network had distinct temporal behaviors, so we then sought to explore whether their real-time patterns carried any representations of each subject’s observable behaviors during the recording period. All subjects were patients confined to a hospital room, undergoing multi-day epilepsy evaluations under video and audio surveillance (**Fig. 1b**). Primary activities that we were able to quantify were thus right and left-hand movements, conversation and (in some cases) eyes open / closed.

Comparing PSD switching patterns of each network to contemporaneous behaviors, we found striking agreements between the switching dynamics of specific networks and behavioral state dynamics (**Fig. 4a,f**). Similarly, correlating behavioral state dynamics with PSD switching dynamics of individual channels revealed coordinated, distributed networks across diverse brain regions (**Fig. 4b,c/g,h**). In both cases, the anatomical locations of electrodes whose PSD switching properties correlated with behavioral state transitions spanned both functional areas that would naturally be expected to participate in the execution of a behavior (e.g. sensory or motor areas), as well as higher cognitive areas such as those involved in motor planning and organization, language comprehension and speech production, object recognition, emotional processing, decision making, working memory, and attention. (**Fig. 3e,k**, **Fig. 4c,h**, **Fig. 5b**, supplementary **Table S4)**.

It is important to note that this analysis examined representations of multiple, simultaneously occurring behavioral streams (e.g. hand movements and conversation) capturing the way in which multiple threads of dynamic network interactions occur in parallel within the real-time brain. Although our data-driven network clustering implies that specific channels belong to a static network structure, networks identified via correlations to specific behaviors suggest that some brain regions can serve overlapping roles in multiple behaviors simultaneously. We interpret these results as revealing real-time, brain-wide representations of network cooperation and communications that depict the parallel flow of information between brain regions involved in the generation and execution of simultaneously occurring spontaneous behaviors.

### Functional exclusivity and stability of networks over days, and during N3 sleep

Our final investigation questioned how such a large percent of electrode channels could be tuned to our relatively simple observable behavioral states (e.g. **Fig. 4a,f**), when our brains possess the capacity to perform and process such a huge repertoire of behaviors and tasks. We attribute the high representation of our observable behaviors to several factors; First, we note that our constrained recording environment likely reduced the dimensionality of behaviors that subjects’ brains were engaged in (**Fig. 1b**). Second, the clinical purpose of electrode placements also generally biases recording locations to auditory and speech processing and sensory and motor regions, which will preferentially encode the execution of relatively simple behaviors. Similarly, we would not expect to find representations of behaviors that engage brain regions that are not sampled by an individual subject’s electrode placements. For example, posterior parietal and occipital cortices were not sampled in this dataset owing to the rarity of seizures arising from these regions, and thus our data would have limited coverage of networks involved in visual function. We also note that additional networks with strong switching patterns were observed, and that these likely correspond to additional dynamic components of behavior or cognition that we have yet to identify, given limitations of the behavioral observations achievable in this study.

Although our results above already suggest somewhat plastic or overlapping roles for each brain region, rather than exclusive and rigid membership of a single network, we sought to further understand the stability of PSD switching network structures by comparing data from the same subject on two different days, and comparing patterns in the awake / alert state to N3 sleep.

Stability of networks in the awake brain from day to day was confirmed (**Fig. 6c,d**, **Fig. 7a,c**, Supplementary **Fig. S8**), underscoring the robust nature of our PSD switching and network classification approach. We note that this property is consistent with the stability of functional networks identified using multi-day fMRI^34^ and iEEG^35^ recordings.

However, as anticipated, network structures were significantly altered during N3 sleep. In particular, channels within strongly synchronized networks in the awake state reduced their intra-network synchrony during N3 sleep, but also became more synchronous overall with the rest of the brain. It is important to note that PSD switching wasn’t absent during N3 sleep (**Fig. 6e**, **Fig. 7d, Fig. S6e** and **Fig. S7e**). However, in networks that were strongly correlated to specific behavioral state transitions (such as conversation) in the awake state, PSD switching during N3 sleep no longer correlated with any of our characterized behaviors. These results are aligned with fMRI BOLD studies observing a “breakdown” in functional connectivity during N3 sleep^36–38^.

It is well known that there can be different drivers of cortical activity during sleep, for example slow wave sleep, which is present during the N3 sleep state, and is associated with network synchrony^39,40^. Our focus on broadband activity between 2-55 Hz, and our use of a 2 s moving window, may have obscured direct observation of slow waves. However, the distinctly different PSDs observed during N3 sleep across recordings sites (**Fig. 6c** (iv)**, e** (iv), **Fig. 7a**, **Fig. S6 c** (iv)**, e** (iv) and **Fig. S7 c** (iv)**, e** (iv)) indicate a state-dependent change in underlying neuronal activity, consistent with both fMRI^41^ and iEEG studies^42^. Our results clearly demonstrate that in the context of changes in state-dependent neural dynamics, the structure of functional networks is not ‘hard-wired’, but exhibits state-dependent plasticity.

### ‘Fast-switching’ channels

The majority of our analysis focused on groups of channels whose PSD switching vectors were strongly correlated to each other. However, in all subjects we found ‘fast-switching’ channels whose PSD switching appeared to be relatively rapid and noisy, and was not strongly correlated to other channels in the awake state. These channels were generally grouped together by hierarchical clustering, but exhibited low intra-network correlation (e.g. **Fig. 7**, FCN4). Surprisingly, these channels became markedly more synchronous with each other, and the rest of the brain, during N3 sleep (**Fig. 7e,g**). Across subjects, these channels were also generally located in similar anatomical areas such as hippocampus, parahippocampal gyrus, amygdala, fusiform gyrus, middle frontal gyrus, orbital part of superior frontal, middle frontal and inferior frontal gyrus (**Fig. 7f** (i)).

More rapidly switching PSDs could simply imply smaller variations between PSD states, and thus unstable binary classification. Alternatively, the dynamics of state switching in these structures may operate on a finer time resolution than the 2 second moving window used in our PSD calculation, or may involve more than 2 states. Given the anatomical locations of these channels, we reason that they might be encoding higher order functions and complicated or faster modulations corresponding to changes in level of arousal or engagement. Loss of information could also result from our exclusion of low delta band (0-2 Hz) activity, which is known to play an important role in coordinating cross-areal interactions related to cognitive function during awake^43,44^. High gamma frequencies were also excluded to reduce noise, but have been associated with internal emotional states during naturalistic behaviors^45^, motor tasks^46^ and language function^47^. Stronger synchrony of fast-switching regions during N3 sleep could relate to slow waves, which typically originate in frontal cortex, propagating along the anterior-posterior axis to mesial temporal lobe and hippocampus^41^, a subset of regions within this fast-switching group.

We also recognized that it was common for this fast-switching group to include channels identified as exhibiting interictal epileptiform discharges (IEDs), which were predominantly observed during sleep. To confirm whether IED events contribute to network correlation properties, in **Fig. S9** we compare inter-channel correlations of PSD switching dynamics, low (1-8 Hz) and high frequency (20-40 Hz) band power during awake and N3 sleep states, separating IED channels from identified networks in each subject. We confirm that IED activity (visible in the high frequency band) is stronger during sleep than awake states, and that IED activity is correlated within IED channels during sleep. However, we find no evidence that IED activity is driving brain-wide synchrony or PSD switching during sleep or awake states. Analysis of the prevalence and spatial distribution of IED channels relative to the fast-switching channels (quantified as ‘mean state duration’) across subjects is shown in **Fig. S10**. Results from demonstrate that IED channels are more commonly (but not exclusively) ‘fast switching’, and that there exist many fast-switching channels that do not exhibit IEDs (**Fig. 7e,g** and **Fig. S10**). Although this analysis is limited by the fact that IED locations were similar across most of the patients in this study, we conclude that fast-switching is likely a common feature of brain regions that have a higher tendency to be sources of epileptiform activity, but found no evidence for a causal relationship between the two phenomena. Nevertheless, we acknowledge that pathological effects of epilepsy could be impacting cognitive function in our subjects^48,49^ and could play a role in some of the network properties observed here^50^.

### The importance of broad-band analysis of network activity

Comparing our results to prior studies, bilaterally symmetric, region-specific differences in time-averaged PSD profiles have been previously observed in the human brain. Our results agree with the spectral distributions characterized during ‘resting state’ in recent electrophysiological studies^27^. However, our PSD-switching results provide a new perspective for seemingly contradictory functional connectivity results obtained using spectral band-limited neural activity.

For example, spectral band-limited studies of human brain electrophysiological activity have reported either no correlation in frequency-specific activities^51^, or connectivity patterns that differ across frequencies^24,52–54^. In some cases, band-limited connectivity patterns have demonstrated overlaps^13,55–57^, and spatial correspondence with fMRI BOLD derived patterns across multiple frequency bands^13,58,59^. Further studies have reported that matching MEG and fMRI correlation structures requires cortical location-dependent frequency selection^60^, or that combining information across multiple frequency bands best represents fMRI-derived connectivity organization^61^. Here, we observed functional networks, defined by synchronous switching of broadband PSDs between at least two states. We found that switching could be synchronous between two regions, even if their PSD patterns differed significantly, and that switching could manifest as both increases or decreases in spectral power in a frequency and region-dependent way (**Fig. 3d,j (ii), Fig. 4d**, **Fig. 5a**, **Fig. 6c-e** (v)). This observed property of brain-wide broadband network activity might explain differences, or absences of correlations, as well as the spatial overlaps, when observations are limited to specific frequency bands, or rely on frequency-dependent correlations as is the case with synchrony metrics generally.

Comparisons to functional connectivity measures derived from resting state fMRI data can be confounded by the complexity of fMRI’s hemodynamic representation of neural activity^6,62,63^. fMRI manifestations of PSD switching could thus be frequency, region or sub-network dependent^60,64^. However, it is anticipated that some level of the neural switching and synchrony dynamics demonstrated here would contribute to detectable fMRI signals^65,66^, and thus could contribute to functional connectivity networks detected in the resting-state and task-active brain^67,68^.

### Physiological interpretations of PSD-switching

Our results build upon our primary observation of PSD switching throughout the real-time brain, a phenomenon that has not been widely described in prior studies. However, we note that dynamic changes in PSD frequency structure have been observed in relation to behavioral state across a range of species and electrophysiological recording approaches: In mice, for example, changes in broadband spectral profiles of local field potentials have been observed during locomotion v/s rest^69^, with evidence of similar shifts in frequency spectrum between different levels of arousal^63^.

A recent modeling study noted that neural activity PSDs do not necessarily denote the presence of oscillatory sources at specific frequencies, but that all time-varying signals will generate rich frequency-domain representations which can change due to a range of effects, from shifts in periodic and quasi-periodic behavior, to changes in the effective impulse response shape of neural responses^70^. In iEEG, electrode channels will be summing signals from a relatively large and diverse population of cells, such that PSD switching could be a simple manifestation of engagement of a different population of neurons or a change in firing patterns, consistent with varying involvement of a brain region in a functionally-relevant behavior. Switching could also reflect the influence of incoming signals to a region, or state-dependent modulation of feedback or resonant activity.

The reliance of most neuroimaging studies on repeated stimuli or tasks, followed by trial averaging may have obscured the presence of real-time PSD switching, while observations at limited locations could lead switching signals to be dismissed as noise or random variability^63^. Our robust results suggest that broadband PSD switching is a widespread and accessible property of the brain-wide network activity underlying real-time processing and naturalistic behaviors in the human brain.

### The value of real-time analysis of representations of naturalistic behaviors

The majority of functional connectivity studies generate spatial representations of networks, but rarely explore interpretation of the real-time dynamics of the activity delineating the network. Conversely, functional studies of the human brain, both using imaging and intracranial recordings, almost always rely on repetitive stimuli or tasks designed to probe specific behaviors or circuits. Trial averaging is typically used to remove the effects of real-time, inter-trials or spontaneous variability. In contrast, by using real-time data, our study uncovered clear neural representations of multiple, concurrent, real-time spontaneous behaviors, across multiple distributed brain networks.

Our analytical approach enables interpretation of multiple threads of neural dynamics, and their relationships to spontaneous behavior in parallel (e.g. left– and right–hand movement, and conversation). Our results provide a brain-wide view of the diverse brain regions whose activity patterns synchronize to decide, generate, execute and process temporally overlapping behavioral streams. We further note that our analysis revealed additional dynamic networks for which we have yet to determine the behavioral / cognitive correlates. We also found compelling ‘disordered’ networks in the awake state which could (along with all other regions) potentially encode higher order PSD switching over faster (or slower) time-scales. Plausibly, additional underlying or even overlapping networks could exist that represent parallel behavioral and cognitive processes across diverse temporal scales^71^. In prospective studies, network functions and properties could be probed via direct or closed-loop interactions with subjects during real-time recordings.

We note that our analysis approach is purely data-driven and uses no predictive modelling. Although several recent investigations have shown that machine learning techniques can decode naturalistic behaviors from multi-day iEEG recordings^45,72,73^, our study exposes the richness, and relative simplicity of information that can be extracted from real-time intracranial recordings. Harnessing real-time PSD-switching, and its synchrony within diverse brain networks could provide improved measurables for machine learning-based analysis, and closed-loop brain computer interfaces. Our results also underscore the importance of studying brain-wide representations of naturalistic behaviors, and provide a new framework for broader interpretation of real-time neural recordings in freely behaving subjects.

## Acknowledgements

The authors acknowledge support from NIH RF1 MH114276 and 1R01NS076628 (Hillman), NIH/NINDS R01 NS084142 and R01 NS110669 (Schevon), and NIH/NIMH R01-MH104606 (Jacobs).

## Author contributions

E.M.C.H and C.A.S. conceived of the study and supervised the project. C.A.S. and H.Z. collected and reviewed the data. E.M.C.H, C.A.S and H.Z developed analysis approaches, interpreted results and prepared the manuscript and figures. A.J.M and A.H.A-M contributed to data acquisition and post-processing. E.M.M, J.J. and M.J.H. contributed to analysis approaches and interpretation of results. C.A.S. and A.J.M reviewed neural data and behavioral annotation and consulted on sleep data analysis. G.M.M, S.A.S and N.A.F. contributed to data collection.

## Ethics statement

Participants provided informed consent for data collected to be utilized for research purposes. All procedures were approved by the Columbia University Irving Medical Center Institutional Review Board.

## Methods

### iEEG and behavioral data acquisition

Ten patients with drug-resistant epilepsy (6 male, 4 female, age 25-55 years, **Table S1**) who underwent stereotactic depth array implantation for seizure localization were included in the study. All participants were hospitalized and monitored in the adult epilepsy monitoring unit (EMU) at New York/Presbyterian Hospital and Columbia University Irving Medical Center. Duration of monitoring ranged from 5 days to 3 weeks, during which anticonvulsant medications were temporarily reduced in order to provoke typical seizures. All data were acquired as part of routine clinical care.

Depth arrays incorporated 8-14 cylindrical recording contacts spaced approximately 5 mm apart (PMT Corp. Chanhassen MN or Ad-Tech Medical Instrumentation Corp, Oak Creek, WI). Simultaneous continuous iEEG and audiovisual recordings were conducted using the Natus Quantum or XLTeK FS128 clinical recording system (Natus Medical Inc, Pleasanton, CA). Electrophysiological data were collected at varying sample rates, specifically 500 Hz (n=1), 512 Hz (n=2) or 1024 Hz (n=7), with +/− 8 mV range and hardware filtering between 0.1 Hz and one fifth of the per channel sampling rate (i.e. 100 – 200 Hz). Depth electrode coverage was tailored to the patients’ suspected epileptogenic areas based on the presurgical evaluation, most commonly in the frontal and temporal lobes but also including parietal coverage in two subjects.

### Inclusion criteria and data selection

Inclusion criteria required participants to have bilateral implants. Power analysis was not performed to determine the sample size due to the limited number of available subjects; however, our sample size is comparable to previous published studies^45,72^. Data were selected from days 2 and 3 after depth array implantation, to avoid lingering effects of general anesthesia and immediate postoperative pain medication, and also effects of anticonvulsant drug reduction and subsequent spontaneous seizures. All behaviors during the selected recording periods were spontaneous and unconstrained. Interictal discharge populations and their fields were identified from clinical procedure reports with confirmation via visual review by HZ with supervision from CS, based on standard clinical criteria. Channels included in these populations were categorized as IED channels for all EEG epochs studied (i.e. the two wake periods and the sleep period).

Data shown in **Fig. 2** includes–5 minutes of ‘resting state’ data from each subject for PSD distribution analysis. Resting state was defined as the patient being awake but at rest in bed without movements, cell phone usage, TV watching, or interaction with others. Interactions before and after the resting period were used to rule out drowsiness/sleep. Two 60-minute recordings were selected during wakefulness, one per day, for each subject **(Fig. 2** and **Fig. 6**, summarized in **Table S2)**. Channels with persistent artifact and time periods with frequent interictal discharges or motion artifacts were excluded from the analysis.

To demonstrate event-related effects in single-trial iEEG data (**Fig. 4e,j**) we identified multiple behavioral state transitions from 24 hours of continuous recordings on day 2. We focused on two naturalistic behaviors displayed by subjects 5 and 6, resulting in 40 events per behavior. Specifically, subject 5 contributed 20 events involving transitions from silence to conversation and 20 events from conversation to silence. Additionally, we extracted 20 events with state transitions from eyes open to closed and 20 events from eyes closed to open for subject 5. Subject 6 contributed 20 events with state transitions from silence to conversation and 20 events from conversation to silence. In addition, we extracted 20 events with state transitions from right hand resting to movement and 20 events from right hand movement to resting for subject 6. Each event had a minimum duration of 5 seconds for each specified behavioral state.

Full night sleep data between days 2 and 3 (supplementary **Fig. S5)** were exported for each individual subject, with a total duration of 73 hours (mean: 7.3 hours/subject, range:4.5-11.3 hours). For subjects 1,4, and 10, data from multiple intermittent sleep periods was also collected due to insufficient continuous sleep duration or frequent sleep interruptions. The selected N3 sleep state is at least 4 hours away from the observed seizure (n=1). The procedure for N3 sleep stage detection is detailed further below.

### Electrode localization

Pre-implant brain magnetic resonance imaging (MRI) and post-implant computed tomography (CT) scans were acquired for each participant. Electrode locations were determined using the Intracranial ELectrode VISualization (iELVis) pipeline^29^. Pre-implant MRI scans were segmented using the FreeSurfer software^74^, and post-implant CT scans were subsequently registered to pre-implant MRI scans through FMRIB Software Library (FSL)^75^. The localization of electrodes within the co-registered CT scans was performed using the BioImage Suite software^76^. An additional expert conducted a visual examination of the results on the co-registered CT and MRI scans to ensure accuracy.

The coordinates of depth electrode contacts for each patient were mapped to the Montreal Neurological Institute (MNI) 305 space via an affine transformation. The anatomical structure of each electrode contact was determined using the Automated anatomical labeling (AAL) atlas^30^ (version SPM12) based on the MNI152 coordinates transformed from MNI305 space. Whether electrodes were within grey or white matter was assessed using the FreeSurfer segmentation results. The MNI152 template was used for brain visualization (Python Nilearn package).

### Spectral analysis

All iEEG channels were average referenced (artifactual channels excluded), notch filtered at 60 Hz and 120 Hz (FIR filter, zero-phase, 2 Hz notch width, Python MNE package, filter.notch filter function), and z-scored. All analyses were programmed in Python.

The sliding-windowed PSD of each channel was calculated using the Welch method (Python Sci-Py, signal.welch function) with a Hann window length of 2 seconds and 50% overlap. This resulted in a sampling rate of 1 second, aligning with the temporal resolution of the behavioral annotation. The PSD within each sliding window was then normalized to a sum of 1. Additionally, the 1/f component of PSD between 2-55 Hz was estimated and removed using the FOOOF algorithm^32^ within each sliding window.

### Spatial PSD distribution analysis

The whitened PSD profiles were time-averaged over the 5-minute resting period for each individual channel, and subsequently averaged over channels within each brain region of interest (**Fig. 2**). The region-averaged PSD profiles were sorted based on their respective peak frequencies, from smallest to greatest. The peak frequency refers to the frequency with the greatest power. Similarly, whitened PSD profiles obtained during the 60-minute awake period on day1 were also time-averaged and region averaged (**Fig. 2**).

### Sliding-windowed PSDs K-means clustering

For each individual channel, transient PSD profiles acquired from each sliding time-window were clustered into two groups using K-means clustering without predefined centroids, with 10 repetitions. This clustering procedure relied solely on iEEG data, independently of any behavioral information. The existence of these two distinct clusters was confirmed for each individual channel using the Duda-Hart test. Additionally, the choice of two clusters was supported by the observation of the lowest silhouette score, indicating a better model fit compared to a greater number of clusters. The two resulting clusters were ranked based on their gamma band power (35-55 Hz) without prior assumptions. Specifically, the cluster with lower power was designated as cluster 0, while the other cluster was denoted as cluster 1. The resulting classification of PSDs results in a ‘PSD switching vector’ for each electrode channel.

We note that this ranking approach was applied to the Day 1 wakefulness and N3 sleep dataset (**Fig. 2 & Fig. 6**). For Day 2 wakefulness, the ranking of the two clusters was based on their similarity compared to each of the clusters characterized during Day 1 wakefulness. This was done to reduce the variation introduced during the 1/f component removal process.

### PSD Network clustering

The PSD switching vectors encode dynamic changes in neural activity over time. To sort PSD switching vectors into correlated groups we used hierarchical clustering. For this analysis, binary PSD switching vectors were first smoothed using a 5-second moving average window and then clustered into networks within each subject using Ward’s method^77^ based on their correlation distances (**Fig. 3**, **Fig. 6** & **Fig. 7**). The correlation distance for a given pair of channels, computed using their PSD switching vectors *u* and *v*, and is defined as:

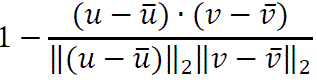

where 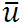 is the mean of the elements of *u* and *x* · *y* is the dot product of *x* and *y*. The optimal number of clusters was determined for each subject based on the silhouette score (supplementary **Fig. S2**). Clustered networks are displayed from top to bottom based on their averaged within-network distance, from highest to lowest. Channels within each network were rearranged based on their mean PSD correlation distance to other channels within network, from the lowest to greatest.

### N3 stage sleep detection

We employed the beta/delta power ratio as a metric for identifying N3 stage sleep in each patient, a method previously validated in human iEEG recordings.^73,78,79^ The spectrogram of each channel was computed using the Welch method (as described above) with a Hann window length of 10 sec and 50% overlap. Following this, we computed the ratio of beta (12-30 Hz) and delta (0.5-4 Hz) band power for each channel, followed by averaging across channels (excluding channels with frequent interictal discharges). Subject-specific thresholds were applied to identify periods corresponding to N3 stage sleep. As a result, an average of 30-minute per subject (range: 20.8-33.3 minutes) was extracted. Subject 4 was excluded due to unstable sleep, and subject 5 was excluded due to the presence of large-scale, frequent epileptiform discharges throughout the full night sleep. For other subjects, time periods with frequent epileptiform discharges within the selected N3 sleep periods were excluded from further analysis.

## Behavioral analysis

### Hand-annotated behavioral labeling

During the initial 60-minute wakeful recording on day 1, we manually annotated spontaneous naturalistic behavioral state transitions in the audio-visual recordings of 10 subjects. These behaviors included conversation engagement, left/right-hand movements, and eyes open vs. eyes closed, which are commonly observed in patients undergoing monitoring in the EMU.

The labeling process relied solely on the audio-visual recordings and was performed independent of the iEEG data. Each behavior was labeled at a one-second temporal resolution, with a value of 0 indicating the absence of the behavior and a value of 1 indicating its presence. “Eyes open (1)” was defined as the presence of at least one eye being observed as open in the video recordings, while “eyes closed (0)” was defined as the closure of both eyes. Blinking episodes lasting less than one second were not considered as instances of eyes closed in this study. Timepoints in which the eyes were neither observed nor obscured were excluded from the analysis. The “conversation (1)” state referred to active language engagement, including both speech production and reception. Conversely, “silence (0)” indicated the absence of any discernible language activity. The study excluded confounding factors such as TV watching and remote conversation occurring outside the monitoring unit. “Hand movement (1)” represented a state in which the subject was observed actively moving their hand or manipulating objects with their hand. In contrast, “hand resting (0)” indicated timepoints during which no such hand movements were observed. Timepoints solely associated with leg movements were excluded. Also, timepoints in which the target hand was obscured or engaged in typing on a cell phone were excluded from the analysis.

To ensure the accuracy of the annotations, the labels were validated by an epilepsy fellow physician with substantial expertise in analyzing the monitoring videos of patients in the EMU. Timepoints affected by motion artifacts, or wire interference were excluded from the study. Instances of low confidence level, such as when certain body parts were obscured or not clearly visible in the camera, were also excluded from the study.

### Audio spectral analysis for ‘conversation’ quantification

Audio information was extracted from audio-visual recordings. The spectrogram of audio data was computed using the Welch method (Python Sci-Py, signal.welch function) with a Hann window length of 2 sec and 50% overlap, resulting in a 1-sec temporal resolution. In **Fig. 6** & **Fig. 7**, we utilized 80 to 400 Hz power that spans human vocal frequencies. The power was summed and normalized to a range of 0 and 1.

### Movement analysis using DeepLabCut

Three body parts, head, left and right hands were tracked using a machine learning pose estimation based on neural networks. For training, 20 frames were extracted and manually labeled for every 2-min video segment (video frame rate=30/sec in 9 subjects and 25/sec in 1 subject). The model was trained for a minimum of 200,000 iterations separately for each recording of day 1 wakefulness, day 2 wakefulness, and N3 sleep state within each subject. In **Fig. 6** & **Fig. 7**, we calculated motion energy for left/right hand between frames and averaged across all frames within every second, resulting in a 1-sec temporal resolution. Outliers that were 3 standard deviations away from the mean were replaced by the mean.

### Behavioral correlation with PSD switching vectors

We evaluated the relationship between PSD switching vectors and spontaneous behaviors using both manual annotation of behavioral state transition and semi-automated extracted behavioral changes. In **Fig. 4** & **Fig. 5**, we first smoothed PSD switching vectors of each individual channel and the annotations of each category of behavioral state transition with a 5-second moving average window and then calculated their relationship using Pearson Correlation. In **Fig. 6** & **Fig. 7** where we semiautomatically extracted audio power, left-/right-hand motion energy from monitoring video, we computed their correlation to PSD switching vectors after smoothing using Spearman Rho. In **Fig. S8**, we mapped the absolute values of correlation coefficients (Spearmanr Rho) found in day 1 and 2 wakefulness onto brain plots.

## Statistical analysis

### PSD profiles comparison in bilateral brain areas

In order to group channels from homologous brain areas (**Fig. 2**), we tested whether channels from the left side are more similar to the right than to those from elsewhere. We investigated the differences in time-averaged PSD profiles characterized from 60-min wakefulness between left and right hemispheres’ homologous brain areas (**Fig. S1a**). For each investigated brain area, we computed the Euclidean distance between the PSDs of channels in the left hemisphere and the following: 1) mean PSD of all channels located in left hemisphere; 2) mean PSD of all channels located in right hemisphere; 3) mean PSDs from each of the other brain areas. We compared the distribution of distances between unilateral and contralateral groups (case 1 and 2), as well as between the contralateral group and cross-ROI group (case 2 and 3). Our null hypotheses were: 1) the mean distance from unilateral group is not smaller than the contralateral group; 2) the mean distance from contralateral group is not smaller than the cross-ROI group. For each null hypothesis, we performed a one-sided Mann-Whitney U test. The analysis was repeated for every investigated brain area, and the results were adjusted for multiple comparisons using the Holm-Sidak method with a significance level set at alpha=0.05. Brain areas with less than 10 channels in either left or right side were excluded from the analysis.

### PSD profiles comparison within network

In **Fig. S2d**, we examined the differences in the time-averaged PSD profiles across channels within the same network. Individual channel’s PSDs within each network were clustered into two groups using hierarchical clustering with the Ward method based on correlation distances. The existence of two clusters was confirmed for each network using the Duda-Hart test. We did not assess the optimal number of clusters as our focus was on the two most distinct clusters. For each network, we assessed the proportion of channels exhibiting different PSDs to the rest of the channels within the network by computing the ratio of the size of smaller cluster compared to the total number of channels.

### PSD variation comparisons across datasets

In **Fig. 2**, we tested whether the transient PSD profiles from the 60min awake period demonstrate more variation than those from the 5min resting period. Our null hypothesis was that the mean PSD variation from 5-min resting period is not smaller than that observed from 60-min wakefulness. For each individual channel, the variation in PSDs is evaluated by the correlation distance between each sliding-windowed PSD profile to the time-averaged PSD profile within each dataset (**Fig. S1b**). At the group level, we performed a one-sided Wilcoxon signed-rank test to compare the distribution of PSD variation between the two datasets.

### PSD profile comparison across clusters

We tested whether the PSD profiles are different between the two K-means clusters, or between annotated behavioral states. In **Fig. 4**, we tested which frequency exhibits a statistically significant difference in power distribution between annotated behavioral states (timepoints with label 0 vs label 1). Out hypothesis was that there are no differences in the mean power across annotated behavioral states. For each frequency, we performed a two-sided Mann-Whitney U test to compare the power distribution across the two states, with corrections applied for multiple comparisons. This analysis was repeated for each investigated channel. In **Fig. 5**, a similar analysis was conducted to test for power differences across K-means clusters. In **Fig. 3** and **Fig. 6** where individual channel examples were presented, we performed a two-sided Mann-Whitney U test to compare the distribution of power centered at their respective dominant frequencies across the two K-means clusters. The dominant frequency is defined as the frequency that explains the most variance in the sliding-windowed whitened PSD profiles.

### Functional selectivity across networks

In **Fig. 5**, we tested whether channels within the identified responsive networks demonstrate a higher level of correlation to the investigated behavioral state change than channels from other networks. For each subject, we first identified channels with a correlation coefficient greater than 0.5 to the investigated behavioral state change. We then identified the network that contains the most of these channels and defined it as the responsive network. Within each subject, we performed a one-sided Mann-Whitney U test to compare the mean of correlation coefficients between channels within the responsive network and channels from other networks. The results were corrected for multiple comparisons across subjects. This analysis was repeated for each category of behavioral state changes.

### PSD profile comparison across datasets

In **Fig. 7a**, we tested whether the PSD profiles characterized from day 2 wakefulness or N3 sleep are more similar to those initially characterized from day 1 awake period. For each individual channel, the similarity of their cluster-averaged PSD profiles across datasets was assessed using the Pearson correlation. Our null hypothesis was that the mean correlation coefficients (correlated to day 1 wakefulness) from day 2 are not greater than those from the N3 sleep state. At the group level, we compared the distribution of similarity between day 2 wakefulness and N3 sleep for each PSD cluster using the one-sided Mann-Whitney U test.

### Correlation matrix comparison across datasets

In **Fig. 7c**, we tested whether the correlation matrices observed from day 2 wakefulness or N3 sleep are more similar to that observed from day 1 awake period. We first sorted PSD switching vectors from day2 wakefulness and N3 sleep based on the network organization characterized from day 1 wakefulness and then computed the correlation matrices. The upper triangular part of each correlation matrix was then vectorized and the similarity across datasets were assessed using the Pearson correlation. Our null hypothesis was that the mean correlation coefficients from day 2 wakefulness are not greater than those from the N3 sleep state. At the group level, the distribution of similarity between day 2 wakefulness and N3 sleep was compared using the one-sided Wilcoxon signed-rank test.

### Comparison of intra-network and inter-network distance

We tested the differences in the distribution of intra-network distance and inter-network distance between wake and sleep. We first assigned individual channel to networks that characterized from day 1 wakefulness. For each individual channel, we computed their averaged correlation distance with channels within the same network, as well as the distance with channels elsewhere. In **Fig. 7f and g**, for each individual channel, we adjusted the inter– and intra-network distance by dividing them by the grand mean of intra-network distance across all channels from the same subject. This adjustment allows for comparison of values across datasets that may have different baseline level of correlation strength. Our null hypotheses were as follows: 1) the variance of adjusted intra-network distance from day 1 wakefulness is not greater than that observed from the N3 sleep state; 2) mean of adjusted inter-network distance from day 1 is not greater than that observed from the N3 sleep state. For null hypothesis 1, we performed a one-sided Wilcoxon signed-rank test to compare the mean of adjusted inter-network distance across the two states. For null hypothesis 2, we performed a one-sided F-test to compare the variance of adjusted distance across datasets. In **Fig. 7h and i**, we mapped the adjusted intra-network distance and their net change between day 1 wakefulness and N3 sleep to brain areas. Brain areas defined by the AAL atlas with less than 10 channels were excluded from the analysis.

## References

1. van den Heuvel, M. P. & Hulshoff Pol, H. E. Exploring the brain network: A review on resting-state fMRI functional connectivity. Eur. Neuropsychopharmacol. 20, 519–534 (2010).

2. Bressler, S. L. & Menon, V. Large-scale brain networks in cognition: emerging methods and principles. Trends Cogn. Sci. 14, 277–290 (2010).

3. Petersen, S. E. & Sporns, O. Brain Networks and Cognitive Architectures. Neuron 88, 207– 219 (2015).

4. M.H. Lee, C.D. Smyser, & J.S. Shimony. Resting-State fMRI: A Review of Methods and Clinical Applications. *Am*. J. Neuroradiol. 34, 1866 (2013).

5. Riedl, V. et al. Metabolic connectivity mapping reveals effective connectivity in the resting human brain. Proc. Natl. Acad. Sci. 113, 428–433 (2016).

6. Hillman, E. M. C. Coupling Mechanism and Significance of the BOLD Signal: A Status Report. Annu. Rev. Neurosci. 37, 161–181 (2014).

7. Allen, E. A. et al. Tracking Whole-Brain Connectivity Dynamics in the Resting State. Cereb. Cortex 24, 663–676 (2014).

8. O’Neill, G. C. et al. Measurement of dynamic task related functional networks using MEG. NeuroImage 146, 667–678 (2017).

9. Liu, Q., Farahibozorg, S., Porcaro, C., Wenderoth, N. & Mantini, D. Detecting large-scale networks in the human brain using high-density electroencephalography. Hum. Brain Mapp. 38, 4631–4643 (2017).

10. Wantzen, P. et al. EEG resting-state functional connectivity: evidence for an imbalance of external/internal information integration in autism. J. Neurodev. Disord. 14, 47 (2022).

11. Sockeel, S., Schwartz, D., Pélégrini-Issac, M. & Benali, H. Large-Scale Functional Networks Identified from Resting-State EEG Using Spatial ICA. PLOS ONE 11, e0146845 (2016).

12. Nir, Y. et al. Interhemispheric correlations of slow spontaneous neuronal fluctuations revealed in human sensory cortex. Nat. Neurosci. 11, 1100–1108 (2008).

13. Aaron Kucyi et al. Intracranial Electrophysiology Reveals Reproducible Intrinsic Functional Connectivity within Human Brain Networks. J. Neurosci. 38, 4230 (2018).

14. Arnulfo, G. et al. Long-range phase synchronization of high-frequency oscillations in human cortex. Nat. Commun. 11, 5363 (2020).

15. Yan, Y. et al. Human cortical networking by probabilistic and frequency-specific coupling. NeuroImage 207, 116363 (2020).

16. Liu, X., Yanagawa, T., Leopold, D. A., Fujii, N. & Duyn, J. H. Robust Long-Range Coordination of Spontaneous Neural Activity in Waking, Sleep and Anesthesia. Cereb. Cortex 25, 2929–2938 (2015).

17. Fox, K. C. R. et al. Intrinsic network architecture predicts the effects elicited by intracranial electrical stimulation of the human brain. *Nat*. Hum. Behav. 4, 1039–1052 (2020).

18. Foster, B. L. & Parvizi, J. Resting oscillations and cross-frequency coupling in the human posteromedial cortex. NeuroImage 60, 384–391 (2012).

19. Brookes, M. J. et al. Investigating the electrophysiological basis of resting state networks using magnetoencephalography. Proc. Natl. Acad. Sci. 108, 16783–16788 (2011).

20. Maldjian, J. A., Davenport, E. M. & Whitlow, C. T. Graph theoretical analysis of resting-state MEG data: Identifying interhemispheric connectivity and the default mode. NeuroImage 96, 88– 94 (2014).

21. Brookes, M. J. et al. Measuring functional connectivity using MEG: Methodology and comparison with fcMRI. NeuroImage 56, 1082–1104 (2011).

22. Sadaghiani, S. & Wirsich, J. Intrinsic connectome organization across temporal scales: New insights from cross-modal approaches. Netw. Neurosci. 4, 1–29 (2020).

23. van Diessen, E. et al. Opportunities and methodological challenges in EEG and MEG resting state functional brain network research. Clin. Neurophysiol. 126, 1468–1481 (2015).

24. Hillebrand, A., Barnes, G. R., Bosboom, J. L., Berendse, H. W. & Stam, C. J. Frequency-dependent functional connectivity within resting-state networks: An atlas-based MEG beamformer solution. NeuroImage 59, 3909–3921 (2012).

25. Jin, S.-H., Seol, J., Kim, J. S. & Chung, C. K. How reliable are the functional connectivity networks of MEG in resting states? J. Neurophysiol. 106, 2888–2895 (2011).

26. Mahjoory, K., Schoffelen, J.-M., Keitel, A. & Gross, J. The frequency gradient of human resting-state brain oscillations follows cortical hierarchies. eLife 9, e53715 (2020).

27. Frauscher, B. et al. Atlas of the normal intracranial electroencephalogram: neurophysiological awake activity in different cortical areas. Brain 141, 1130–1144 (2018).

28. Taylor, P. N. et al. Normative brain mapping of interictal intracranial EEG to localize epileptogenic tissue. Brain 145, 939–949 (2022).

29. Groppe, D. M. et al. iELVis: An open source MATLAB toolbox for localizing and visualizing human intracranial electrode data. J. Neurosci. Methods 281, 40–48 (2017).

30. Tzourio-Mazoyer, N. et al. Automated Anatomical Labeling of Activations in SPM Using a Macroscopic Anatomical Parcellation of the MNI MRI Single-Subject Brain. NeuroImage 15, 273–289 (2002).

31. Mathis, A. et al. DeepLabCut: markerless pose estimation of user-defined body parts with deep learning. Nat. Neurosci. 21, 1281–1289 (2018).

32. Donoghue, T. et al. Parameterizing neural power spectra into periodic and aperiodic components. Nat. Neurosci. 23, 1655–1665 (2020).

33. Khan, Z. U., Martín-Montañez, E. & Baxter, M. G. Visual perception and memory systems: from cortex to medial temporal lobe. Cell. Mol. Life Sci. 68, 1737–1754 (2011).

34. Gratton, C. et al. Functional Brain Networks Are Dominated by Stable Group and Individual Factors, Not Cognitive or Daily Variation. Neuron 98, 439–452.e5 (2018).

35. Chapeton, J. I., Inati, S. K. & Zaghloul, K. A. Stable functional networks exhibit consistent timing in the human brain. Brain 140, 628–640 (2017).

36. Tagliazucchi, E. & Laufs, H. Decoding Wakefulness Levels from Typical fMRI Resting-State Data Reveals Reliable Drifts between Wakefulness and Sleep. Neuron 82, 695–708 (2014).

37. Jobst, B. M. et al. Increased Stability and Breakdown of Brain Effective Connectivity During Slow-Wave Sleep: Mechanistic Insights from Whole-Brain Computational Modelling. Sci. Rep. 7, 4634 (2017).

38. Victor I. Spoormaker et al. Development of a Large-Scale Functional Brain Network during Human Non-Rapid Eye Movement Sleep. J. Neurosci. 30, 11379 (2010).

39. Dang-Vu, T. T. et al. Spontaneous neural activity during human slow wave sleep. Proc. Natl. Acad. Sci. 105, 15160–15165 (2008).

40. Aedo-Jury, F., Schwalm, M., Hamzehpour, L. & Stroh, A. Brain states govern the spatio-temporal dynamics of resting-state functional connectivity. eLife 9, e53186 (2020).

41. Song, C., Boly, M., Tagliazucchi, E., Laufs, H. & Tononi, G. fMRI spectral signatures of sleep. Proc. Natl. Acad. Sci. 119, e2016732119 (2022).

42. Lendner, J. D. et al. An electrophysiological marker of arousal level in humans. eLife 9, e55092 (2020).

43. Nácher, V., Ledberg, A., Deco, G. & Romo, R. Coherent delta-band oscillations between cortical areas correlate with decision making. Proc. Natl. Acad. Sci. 110, 15085–15090 (2013).

44. Harmony, T. The functional significance of delta oscillations in cognitive processing. Front. Integr. Neurosci. 7, (2013).

45. Bijanzadeh, M. et al. Decoding naturalistic affective behaviour from spectro-spatial features in multiday human iEEG. *Nat*. Hum. Behav. 6, 823–836 (2022).

46. Crone, N. E., Miglioretti, D. L., Gordon, B. & Lesser, R. P. Functional mapping of human sensorimotor cortex with electrocorticographic spectral analysis. II. Event-related synchronization in the gamma band. Brain 121, 2301–2315 (1998).

47. Crone, N. E., Boatman, D., Gordon, B. & Hao, L. Induced electrocorticographic gamma activity during auditory perception. Clin. Neurophysiol. 112, 565–582 (2001).

48. Holmes, G. L. & Lenck-Santini, P.-P. Role of interictal epileptiform abnormalities in cognitive impairment. Epilepsy Behav. 8, 504–515 (2006).

49. Holmes, G. L. Cognitive impairment in epilepsy: the role of network abnormalities. Epileptic. Disord. 17, 101–116 (2015).

50. Dahal, P. et al. Interictal epileptiform discharges shape large-scale intercortical communication. Brain 142, 3502–3513 (2019).

51. Deligianni, F., Centeno, M., Carmichael, D. W. & Clayden, J. D. Relating resting-state fMRI and EEG whole-brain connectomes across frequency bands. Front. Neurosci. 8, (2014).

52. Betzel, R. F. et al. Structural, geometric and genetic factors predict interregional brain connectivity patterns probed by electrocorticography. *Nat*. Biomed. Eng. 3, 902–916 (2019).

53. Hipp, J. F., Hawellek, D. J., Corbetta, M., Siegel, M. & Engel, A. K. Large-scale cortical correlation structure of spontaneous oscillatory activity. Nat. Neurosci. 15, 884–890 (2012).

54. Hacker, C. D., Snyder, A. Z., Pahwa, M., Corbetta, M. & Leuthardt, E. C. Frequency-specific electrophysiologic correlates of resting state fMRI networks. NeuroImage 149, 446–457 (2017).

55. Nentwich, M. et al. Functional connectivity of EEG is subject-specific, associated with phenotype, and different from fMRI. NeuroImage 218, 117001 (2020).

56. Parham Mostame & Sepideh Sadaghiani. Oscillation-Based Connectivity Architecture Is Dominated by an Intrinsic Spatial Organization, Not Cognitive State or Frequency. J. Neurosci. 41, 179 (2021).

57. Bassett, D. S., Meyer-Lindenberg, A., Achard, S., Duke, T. & Bullmore, E. Adaptive reconfiguration of fractal small-world human brain functional networks. Proc. Natl. Acad. Sci. 103, 19518–19523 (2006).

58. Wirsich, J. et al. Complementary contributions of concurrent EEG and fMRI connectivity for predicting structural connectivity. NeuroImage 161, 251–260 (2017).

59. Ridley, B. et al. Simultaneous Intracranial EEG-fMRI Shows Inter-Modality Correlation in Time-Resolved Connectivity Within Normal Areas but Not Within Epileptic Regions. Brain Topogr. 30, 639–655 (2017).

60. Hipp, J. F. & Siegel, M. BOLD fMRI Correlation Reflects Frequency-Specific Neuronal Correlation. Curr. Biol. 25, 1368–1374 (2015).

61. Tewarie, P. et al. Predicting haemodynamic networks using electrophysiology: The role of non-linear and cross-frequency interactions. NeuroImage 130, 273–292 (2016).

62. Ma, Y., et al. Resting-state hemodynamics are spatiotemporally coupled to synchronized and symmetric neural activity in excitatory neurons. Proc. Natl. Acad. Sci. 113, E8463–E8471 (2016).

63. Shahsavarani, S. et al. Cortex-wide neural dynamics predict behavioral states and provide a neural basis for resting-state dynamic functional connectivity. Cell Rep. 42, 112527 (2023).

64. Hall, E. L., Robson, S. E., Morris, P. G. & Brookes, M. J. The relationship between MEG and fMRI. Multimodal Data Fusion 102, 80–91 (2014).

65. Stevenson, C. M., Brookes, M. J. & Morris, P. G. β-Band correlates of the fMRI BOLD response. Hum. Brain Mapp. 32, 182–197 (2011).

66. Winterer, G. et al. Complex relationship between BOLD signal and synchronization/desynchronization of human brain MEG oscillations. Hum. Brain Mapp. 28, 805– 816 (2007).

67. Smith, S. M. et al. Correspondence of the brain’s functional architecture during activation and rest. Proc. Natl. Acad. Sci. 106, 13040–13045 (2009).

68. Lipkin, B. et al. Probabilistic atlas for the language network based on precision fMRI data from >800 individuals. Sci. Data 9, 529 (2022).

69. Vinck, M., Batista-Brito, R., Knoblich, U. & Cardin, J. A. Arousal and Locomotion Make Distinct Contributions to Cortical Activity Patterns and Visual Encoding. Neuron 86, 740–754 (2015).

70. Perrenoud, Q. & Cardin, J. A. Beyond rhythm – a framework for understanding the frequency spectrum of neural activity. Front. Syst. Neurosci. 17, (2023).

71. Vidaurre, D. et al. Spontaneous cortical activity transiently organises into frequency specific phase-coupling networks. Nat. Commun. 9, 2987 (2018).

72. Wang, N. X. R., Olson, J. D., Ojemann, J. G., Rao, R. P. N. & Brunton, B. W. Unsupervised Decoding of Long-Term, Naturalistic Human Neural Recordings with Automated Video and Audio Annotations. Front. Hum. Neurosci. 10, (2016).

73. Kremen, V. et al. Automated unsupervised behavioral state classification using intracranial electrophysiology. J. Neural Eng. 16, 026004 (2019).

74. Fischl, B. et al. Whole Brain Segmentation: Automated Labeling of Neuroanatomical Structures in the Human Brain. Neuron 33, 341–355 (2002).

75. Jenkinson, M. & Smith, S. A global optimisation method for robust affine registration of brain images. Med. Image Anal. 5, 143–156 (2001).

76. Joshi, A. et al. Unified Framework for Development, Deployment and Robust Testing of Neuroimaging Algorithms. Neuroinformatics 9, 69–84 (2011).

77. Ward Jr.., J. H.. Hierarchical Grouping to Optimize an Objective Function. J. Am. Stat. Assoc. 58, 236–244 (1963).

78. Krakovská, A. & Mezeiová, K. Automatic sleep scoring: A search for an optimal combination of measures. Artif. Intell. Med. 53, 25–33 (2011).

79. Reed, C. M. et al. Automatic detection of periods of slow wave sleep based on intracranial depth electrode recordings. J. Neurosci. Methods 282, 1–8 (2017).

