## Supplementary Tables and Figures for "Spectral-switching analysis reveals real-time neuronal network representations of concurrent spontaneous naturalistic behaviors in human brain"

### Supplementary information

#### Supplementary Tables:

| | |
|---|---|
| **Table S1** | Implant summary. |
| **Table S2** | Data summary. |
| **Table S3** | List of anatomical regions (AAL atlas) and their related function. |
| **Table S4** | Behavioral state transition responsive channels and locations. |

#### Supplementary Figures:

| | |
|---|---|
| **Fig. S1** | Bilateral symmetry test for time-averaged PSD profiles and spatial mapping of variation in transient PSDs. |
| **Fig. S2** | Number of clusters in sliding-windowed PSD profiles and functional networks. |
| **Fig. S3** | Networks in individual subjects characterized from day 1 wakefulness. |
| **Fig. S4** | Optimal time lag between PSD switching and behavioral annotation. |
| **Fig. S5** | Full night sleep data and selected N3 sleep state Beta/delta power ratio across full night sleep for each subject. |
| **Fig. S6** | Comparing network dynamic results across days and sleep state (subject 7, as shown in Fig. 6). |
| **Fig. S7** | Comparing network dynamic results across days and sleep state (subject 1, as shown in Fig. 6). |
| **Fig. S8** | Correlation between PSD switching vectors to extracted left/right hand motion energy and audio power from simultaneous audio-visual recordings during wakefulness on day 1 and 2. |
| **Fig. S9** | Correlation map and inter-network distance changes between normal channels and channels with frequent interictal epileptiform discharges (IED) during sleep. |
| **Fig. S10** | Mean state duration brain mapping and IED channel distributions. |

| Subject | Implant location | Epileptogenic zone | #Arrays/ Electrodes | Imaging findings |
|---|---|---|---|---|
| 1 (25/M) | R F-T and LT | R hippocampus and basal temporal region | 12/126 | B/L asymmetric posterior periventricular nodular heterotopias |
| 2 (35/F) | L F-T and RT | L mesial temporal (or L hippocampus) | 11/122 | Nonlesional |
| 3 (39/M) | B/L F-T | R mesial temporal, L OF | 16/174 | R MTS |
| 4 (30/M) | L F-T, R F | L hippocampus and insula | 18/202 | Nonlesional |
| 5 (55/F) | R F-T, L T | R posterior-lateral temporal region | 13/134 | Nonlesional |
| 6 (30/M) | B/L F-T-P | R mesial posterior frontal region | 14/166 | Nonlesional |
| 7 (35/M) | B/L F-T | R posterior cingulate | 17/190 | Nonlesional |
| 8 (44/F) | B/L F-P | L anterior cingulate | 11/130 | Nonlesional |
| 9 (30/F) | B/L F-T | B/L mesial temporal | 13/144 | Possible R MTS |
| 10 (32/M) | B/L T | B/L mesial temporal | 17/224 | Nonlesional |

**Table S1 Implant summary.** Summary of clinical data for 10 subjects (as shown in **Fig. 1**), including demographic information, the location covered by the implant, the location of inferred epileptogenic zone, number of electrodes and total contacts, and pathology results. All patients have had implants in both left and right hemispheres with different coverage, determined by clinical needs. R=right; L=left; B/L = bilateral; F=frontal; OF=orbitofrontal; T=temporal; P=parietal; MTS=mesial temporal sclerosis; FCD=focal cortical dysplasia.

| Subject | Resting state (sec) | Day 1 wakefulness (sec) | Day 2 wakefulness (sec) | Total length of sleep data (hour) | Selected N3 sleep data (sec) |
|---|---|---|---|---|---|
| Subject1 | 270 | 3602 | 3602 | 11.2 | 2000 |
| Subject2 | 301.9 | 3602.6 | 3602.6 | 5.6 | 2000 |
| Subject3 | 301.9 | 3601.5 | 3603.6 | 6 | 2000 |
| Subject4 | 298.9 | 4771.5 | 3602.5 | 11.3 | N/A |
| Subject5 | 238.4 | 4203.1 | 3603.6 | 6.4 | N/A |
| Subject6 | 302.8 | 3612.3 | 3601.6 | 6.5 | 1900 |
| Subject7 | 302.8 | 3631.9 | 3602.6 | 4.5 | 2000 |
| Subject8 | 304.8 | 3665.6 | 3601.6 | 6.9 | 2000 |
| Subject9 | 362.3 | 4021.5 | 3602.6 | 6.4 | 2000 |
| Subject10 | 271.5 | 3600.6 | 3601.6 | 8.46 | 1250 |

**Table S2 Data summary.** Summary of the data for 10 subjects, including resting state, two awake period from two consecutive days, full night sleep and selected N3 sleep state (as shown in **Fig. 2**, **Fig. 3** and **Fig. 6**).

| AAL label | Abbreviation | Anatomical region | Related function |
|---|---|---|---|
| Precentral_L | PreCG | Precentral gyrus | Contains primary motor cortex; responsible for the control of voluntary movement on the body's contralateral side |
| Precentral_R | PreCG | Precentral gyrus | Contains primary motor cortex; responsible for the control of voluntary movement on the body's contralateral side |
| Frontal_Sup_L | SFG | Superior frontal gyrus, dorsolateral | Working memory, spatial processing (dominant side) |
| Frontal_Sup_R | SFG | Superior frontal gyrus, dorsolateral | Impulse control; inhibitory control and motor urgency (nondominant side) |
| Frontal_Sup_Orb_L | SFGorb | Superior frontal gyrus, orbital part | Involved in reward and decision making (positive reward); emotional processing and regulation |
| Frontal_Sup_Orb_R | SFGorb | Superior frontal gyrus, orbital part | Involved in reward and decision making (positive reward); emotional processing and regulation |
| Frontal_Mid_L | MFG | Middle frontal gyrus | Development of literacy (dominant side) |
| Frontal_Mid_R | MFG | Middle frontal gyrus | Responsible for numeracy (nondominant side) |
| Frontal_Mid_Orb_L | MFGorb | Middle frontal gyrus, orbital part | Involved in reward and decision making (abstract rewards); emotional processing and regulation |
| Frontal_Mid_Orb_R | MFGorb | Middle frontal gryus, orbital part | Involved in reward and decision making (abstract rewards); emotional processing and regulation |
| Frontal_Inf_Oper_L | IFGoperc | Inferior frontal gyrus, opercular part | Involved in language processing, comprehension and production; tongue and mouth movement during articulation; tone recognition in language (dominant side) |
| Frontal_Inf_Oper_R | IFGoperc | Inferior frontal gyrus, opercular part | Involved in language processing, comprehension and production; tone recognition in language |
| Frontal_Inf_Tri_L | IFGtriang | Inferior frontal gyrus, triangular part | Involved in semantic processing; language comprehension and translation; (dominant side) |
| Frontal_Inf_Tri_R | IFGtriang | Inferior frontal gyrus, triangular part | Involved in semantic processing; language comprehension and translation |
| Frontal_Inf_Orb_L | IFGorb | Inferior frontal gyrus, orbital part | involved in language processing and comprehension; emotional recognition |
| Frontal_Inf_Orb_R | IFGorb | Inferior frontal gyrus, orbital part | involved in language processing and comprehension; emotional recognition |
| Rolandic_Oper_L | ROL | Rolandic operculum | Involved in sensory, motor, autonomic, cognitive processing and language |
| Rolandic_Oper_R | ROL | Rolandic operculum | Involved in sensory, motor, autonomic, cognitive processing and language |
| Supp_Motor_Area_L | SMA | Supplementary motor area | Control of postural stability; coordinating temporal sequences of actions, bimanual coordination; inhibition of internally generated movement |
| Supp_Motor_Area_R | SMA | Supplementary motor area | Control of postural stability; coordinating temporal sequences of actions, bimanual |

| | | | |
|---|---|---|---|
| | | | coordination; inhibition of internally generated movement |
| Olfactory_L | OLF | Olfactory cortex | Response to odor; odor discrimination and odor memory |
| Olfactory_R | OLF | Olfactory cortex | Response to odor; odor discrimination and odor memory |
| Frontal_Sup_Medial_L | SFGmedial | Superior frontal gyrus, medial | Involved in motor and cognitive control |
| Frontal_Sup_Medial_R | SFGmedial | Superior frontal gyrus, medial | Involved in motor and cognitive control |
| Frontal_Med_Orb_L | SFGmedialorb | Superior frontal gyrus, medial orbital | involved in emotional processing, decision-making, memory, self-perception, social cognition |
| Frontal_Med_Orb_R | SFGmedialorb | Superior frontal gyrus, medial orbital | involved in emotional processing, decision-making, memory, self-perception, social cognition |
| Rectus_L | REC | Gyrus rectus | Unclear (may participate emotional and cognitive processes) |
| Rectus_R | REC | Gyrus rectus | Unclear (may participate emotional and cognitive processes) |
| Insula_L | INS | Insula | Sensorimotor processing; socio-emotional processing; cognitive processing |
| Insula_R | INS | Insula | Sensorimotor processing; socio-emotional processing; cognitive processing |
| Cingulum_Ant_L | ACC | Anterior cingulate & paracingulate gyri | Processing emotions and regulating the endocrine and autonomic responses to emotions |
| Cingulum_Ant_R | ACC | Anterior cingulate & paracingulate gyri | Processing emotions and regulating the endocrine and autonomic responses to emotions |
| Cingulum_Mid_L | MCC | Middle cingulate & paracingulate gyri | Cognitive processing, involving reward-based decision making |
| Cingulum_Mid_R | MCC | Middle cingulate & paracingulate gyri | Cognitive processing, involving reward-based decision making |
| Cingulum_Post_L | PCC | Posterior cingulate gyrus | Visuospatial orientation |
| Cingulum_Post_R | PCC | Posterior cingulate gyrus | Visuospatial orientation |
| Hippocampus_L | HIP | Hippocampus | Memory consolidation; decision-making; spatial navigation |
| Hippocampus_R | HIP | Hippocampus | Memory consolidation; decision-making; spatial navigation |
| ParaHippocampal_L | PHG | Parahippocampal gyrus | Memory encoding and retrieval; spatial navigation |
| ParaHippocampal_R | PHG | Parahippocampal gyrus | Memory encoding and retrieval; spatial navigation |
| Amygdala_L | AMYG | Amygdala | Responsible for the control of emotions and behavior besides memory formation |
| Amygdala_R | AMYG | Amygdala | Responsible for the control of emotions and behavior besides memory formation |
| Calcarine_L | CAL | Calcarine fissure and surrounding cortex | Associated with visual cortex, near primary visual cortex; involved in central and peripheral visual field |

| | | | |
|---|---|---|---|
| Calcarine_R | CAL | Calcarine fissure and surrounding cortex | Associated with visual cortex, near primary visual cortex; involved in central and peripheral visual field |
| Cuneus_L | CUN | Cuneus | Basic visual processing |
| Cuneus_R | CUN | Cuneus | Basic visual processing |
| Lingual_L | LING | Lingual gyrus | Processing visual information; encoding visual memory; identification of words; |
| Lingual_R | LING | Lingual gyrus | Processing visual information; encoding visual memory; identification of words; |
| Occipital_Sup_L | SOG | Superior occipital gyrus | Involved in visual processing; face and object recognition; |
| Occipital_Sup_R | SOG | Superior occipital gyrus | Involved in visual processing; face and object recognition; |
| Occipital_Mid_L | MOG | Middle occipital gyrus | Involved in visual processing; visual perception; object and face recognition; spatial information perception and processing |
| Occipital_Mid_R | MOG | Middle occipital gyrus | Involved in visual processing; visual perception; object and face recognition; spatial information perception and processing |
| Occipital_Inf_L | IOG | Inferior occipital gyrus | Related to visual function of processing faces |
| Occipital_Inf_R | IOG | Inferior occipital gyrus | Related to visual function of processing faces |
| Fusiform_L | FFG | Fusiform gyrus | Face perception; object recognition; reading |
| Fusiform_R | FFG | Fusiform gyrus | Face perception; object recognition; reading |
| Postcentral_L | PoCG | Postcentral gyrus | Perceives somatic sensations from the body; Contains primary somatosensory cortex; |
| Postcentral_R | PoCG | Postcentral gyrus | Perceives somatic sensations from the body; Contains primary somatosensory cortex; |
| Parietal_Sup_L | SPG | Superior parietal gyrus | Involved in spatial cognition and visual perception; somatosensory processing; and other cognitive functions |
| Parietal_Sup_R | SPG | Superior parietal gyrus | Involved in spatial cognition and visual perception; somatosensory processing; and other cognitive functions |
| Parietal_Inf_L | IPL | Inferior parietal gyrus | Involved in perception of emotions; sensory integration; visual-spatial processing; involved in many cognitive functions |
| Parietal_Inf_R | IPL | Inferior parietal gyrus | Involved in perception of emotions; sensory integration; visual-spatial processing; involved in many cognitive functions |
| SupraMarginal_L | SMG | Supramarginal gyrus | Interprets tactile sensory information; involved in perception of space and limb's location |
| SupraMarginal_R | SMG | Supramarginal gyrus | Interprets tactile sensory information; involved in perception of space and limb's location |
| Angular_L | ANG | Angular gyrus | Involved in semantic processing; word reading and comprehension; number |

| | | | |
|---|---|---|---|
| | | | processing; attention and spatial cognition; memory retrieval; conflict resolution |
| Angular_R | ANG | Angular gyrus | Involved in semantic processing; word reading and comprehension; number processing; attention and spatial cognition; memory retrieval; conflict resolution |
| Precuneus_L | PCUN | Precuneus | Involved in memory-related tasks; self-consciousness; visuospatial function |
| Precuneus_R | PCUN | Precuneus | Involved in memory-related tasks; self-consciousness; visuospatial function |
| Paracentral_Lobule_L | PCL | Paracentral lobule | Motor and sensory function for contralateral lower limbs |
| Paracentral_Lobule_R | PCL | Paracentral lobule | Motor and sensory function for contralateral lower limbs |
| Caudate_L | CAU | Caudate nucleus | Planning the execution of movement; also related to cognitive functions such as learning, memory, emotion, rewarding etc. |
| Caudate_R | CAU | Caudate nucleus | Planning the execution of movement; also related to cognitive functions such as learning, memory, emotion, rewarding etc. |
| Putamen_L | PUT | Lenticular nucleus, Putamen | Involved in learning and motor control; speech and language function; rewarding and addiction |
| Putamen_R | PUT | Lenticular nucleus, Putamen | Involved in learning and motor control; speech and language function; rewarding and addiction |
| Pallidum_L | PAL | Lenticular nucleus, Pallidum | Control of conscious and proprioceptive movements |
| Pallidum_R | PAL | Lenticular nucleus, Pallidum | Control of conscious and proprioceptive movements |
| Thalamus_L | THA | Thalamus | Relaying sensory and motor signal, regulation of consciousness and alternes |
| Thalamus_R | THA | Thalamus | Relaying sensory and motor signal, regulation of consciousness and alternes |
| Heschl_L | HES | Heschl's gyrus | Include primary auditory area; related to auditory perception and processing; left is the dominant side |
| Heschl_R | HES | Heschl's gyrus | Include primary auditory area; related to auditory perception and processing; |
| Temporal_Sup_L | STG | Superior temporal gyrus | Involved in sound processing; auditory processing; language comprehension; |
| Temporal_Sup_R | STG | Superior temporal gyrus | Involved in sound processing; auditory processing; language comprehension; |
| Temporal_Pole_Sup_L | TPOsup | Temporal pole: superior temporal gyrus | Involved in visual cognition function; language and semantic processing; socio-emotional function; autobiographic memory function |
| Temporal_Pole_Sup_R | TPOsup | Temporal pole: superior temporal gyrus | Involved in visual cognition function; language and semantic processing; socio-emotional function; autobiographic memory function |
| Temporal_Mid_L | MTG | Middle temporal gyrus | Involved in audio-visual recognition; language processing and semantic memory |

| | | | |
|---|---|---|---|
| Temporal_Mid_R | MTG | Middle temporal gyrus | Involved in audio-visual recognition; language processing and semantic memory |
| Temporal_Pole_Mid_L | TPOmid | Temporal pole: middle temporal gyrus | Involved in visual cognition function; language and semantic processing; socio-emotional function; autobiographic memory function |
| Temporal_Pole_Mid_R | TPOmid | Temporal pole: middle temporal gyrus | Involved in visual cognition function; language and semantic processing; socio-emotional function; autobiographic memory function |
| Temporal_Inf_L | ITG | Inferior temporal gyrus | Involved in visual processing; object recognition; semantic recognition and processing |
| Temporal_Inf_R | ITG | Inferior temporal gyrus | Involved in visual processing; object recognition; semantic recognition and processing |
| Cerebelum_Crus1_L | CERCRU1 | Crus I of cerebellar hemisphere | |
| Cerebelum_Crus1_R | CERCRU1 | Crus I of cerebellar hemisphere | |
| Cerebelum_Crus2_L | CERCRU2 | Crus II of cerebellar hemisphere | |
| Cerebelum_Crus2_R | CERCRU2 | Crus II of cerebellar hemisphere | |
| Cerebelum_3_L | CER3 | Lobule III of cerebellar hemisphere | |
| Cerebelum_3_R | CER3 | Lobule III of cerebellar hemisphere | |
| Cerebelum_4_5_L | CER4_5 | Lobule IV,V of cerebellar hemisphere | |
| Cerebelum_4_5_R | CER4_5 | Lobule IV,V of cerebellar hemisphere | |
| Cerebelum_6_L | CER6 | Lobule VI of cerebellar hemisphere | |
| Cerebelum_6_R | CER6 | Lobule VI of cerebellar hemisphere | |
| Cerebelum_7b_L | CER7b | Lobule VIIB of cerebellar hemisphere | |
| Cerebelum_7b_R | CER7b | Lobule VIIB of cerebellar hemisphere | |
| Cerebelum_8_L | CER8 | Lobule VIII of cerebellar hemisphere | |
| Cerebelum_8_R | CER8 | Lobule VIII of cerebellar hemisphere | |

| | | |
|---|---|---|
| Cerebelum_9_L | CER9 | Lobule IX of cerebellar hemisphere |
| Cerebelum_9_R | CER9 | Lobule IX of cerebellar hemisphere |
| Cerebelum_10_L | CER10 | Lobule X of cerebellar hemisphere |
| Cerebelum_10_R | CER10 | Lobule X of cerebellar hemisphere |
| Vermis_1_2 | VER1_2 | Lobule I,II of vermis |
| Vermis_3 | VER3 | Lobule III of vermis |
| Vermis_4_5 | VER4_5 | Lobule IV,V of vermis |
| Vermis_6 | VER6 | Lobule VI of vermis |
| Vermis_7 | VER7 | Lobule VII of vermis |
| Vermis_8 | VER8 | Lobule VIII of vermis |
| Vermis_9 | VER9 | Lobule IX of vermis |
| Vermis_10 | VER10 | Lobule X of vermis |
| Unknown | NA | Unknown |

**Table S3 List of anatomical regions (AAL atlas) and their related function.** List of regions of interest defined by Automated Anatomical Labeling (AAL) atlas[30], including their respective AAL label, abbreviation and associated function (as shown in **Fig. 2** and **Fig. 5**).

| Behavioral category | Total number of responsive channels across all subjects | Responsive regions |
|---|---|---|
| Eyes open/closed | 73 | Middle temporal gyrus (L/R, n=6 in 3 subj);<br>Hippocampus (L/R, n=6 in 3 subj);<br>Middle frontal gyrus (R, n=3 in 1 subj);<br>Anterior cingulum (L/R, n= 3 in 1 subj);<br>Others (n=55 in 3 subj) |
| Right hand movement/resting | 125 | Supplementary motor area (L/R, n=19 in 3 subj);<br>Precentral gyrus (L/R, n=15 in 3 subj);<br>Postcentral gyrus (L, n=14 in 2 subj);<br>Superior frontal gyrus (L/R, n=13 in 5 subj);<br>Middle cingulum (L/R, n=12 in 5 subj);<br>Middle frontal gyrus (L/R, n=8 in 1 subj)<br>Anterior cingulum (L/R, n=6 in 1 subj);<br>Others (n=38 in 6 subj) |
| Left hand movement/resting | 114 | Middle cingulum (L/R, n=17 in 4 subj);<br>Precentral gyrus (L/R, n=13 in 2 subj);<br>Postcentral gyrus (L/R, n=13 in 2 subj);<br>Supplementary motor area (L/R, n=12 in 3 subj);<br>Middle frontal gyrus (L/R, n=9 in 1 subj)<br>Anterior cingulum (L/R, n=7 in 1 subj);<br>Superior frontal gyrus (L/R, n=6 in 3 subj);<br>Precuneus (L, n=4 in 1 subj)<br>Supramarginal gyrus (R, n=3 in 1 subj)<br>Paracentral lobule (L, n=3 in 1 subj)<br>Others (n=27 in 4 subj) |
| Conversation/silence | 83 | Superior temporal gyrus (L/R, n=44 in 8 subj);<br>Middle temporal gyrus (L/R, n=12 in 2 subj);<br>Heschl's gyrus (L/R, n=7 in 3 subj);<br>Rolandic operculum (L, n=4 in 1 subj);<br>Postcentral gyrus (L, n=4 in 1 subj)<br>Others (n=12 in 5 subj) |

**Table S4 Behavioral state transition responsive channels and locations.** List of responsive channels and their corresponding regions to characterized behavioral state transitions across all subjects (as shown in **Fig. 5**), including eye open/closed, right- and left-hand movement/resting and conversation/silence. Responsive channels are defined as those with a Pearson correlation coefficient greater than 0.5 (out of a total of 1612). All responsive channels demonstrate statistically significant differences in their PSDs between two clusters ($p \leq 0.05$, Mann-Whitney U test, two-sided, FWER corrected**).** subj=Subject.

#### Supplemental Figures

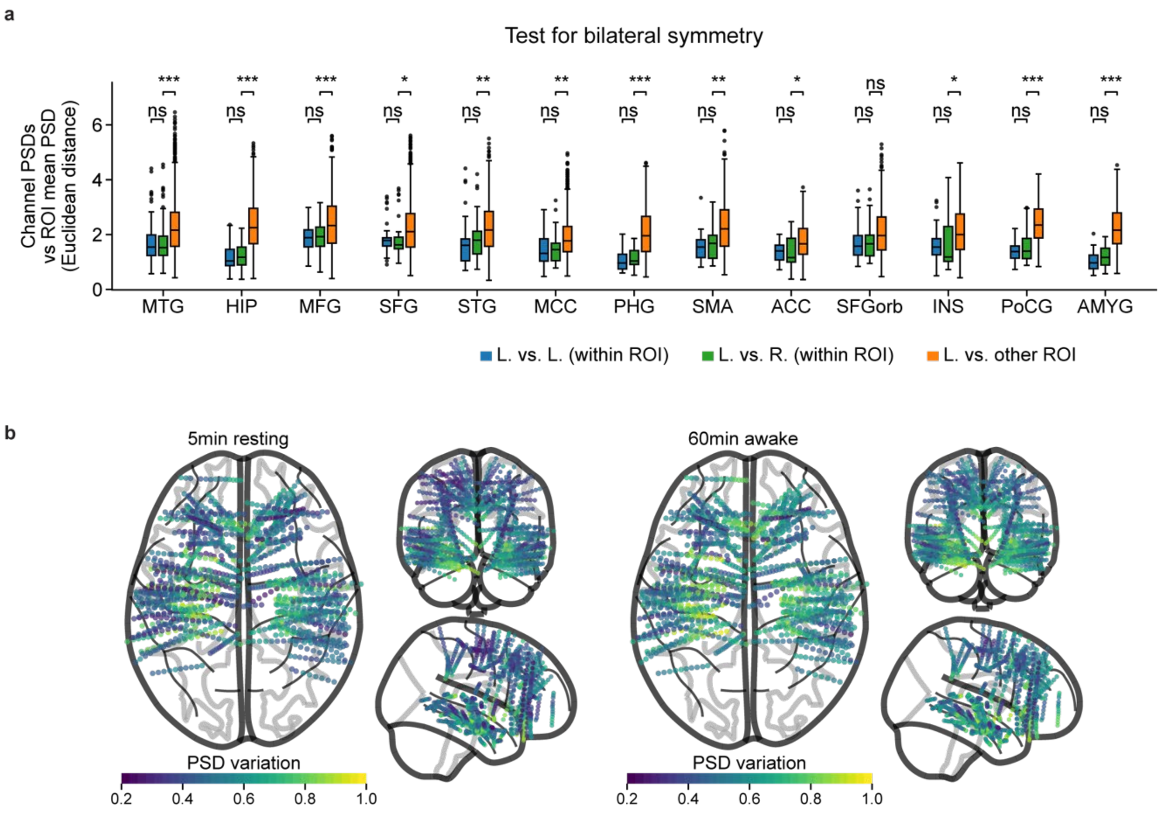

**Fig. S1 Bilateral symmetry test for time-averaged PSD profiles and spatial mapping of variation in transient PSDs. a**, Distribution of individual channel's PSD distance from the mean PSD of each brain area (as shown in **Fig. 2**). No statistically significant differences were found in the mean distance of unilateral and contralateral groups within each ROI ($p>0.05$, Mann-Whitney U test, one-sided, FWER corrected). The mean distance of contralateral groups within the same ROI is significantly smaller than the distance between channels within each respective ROI versus elsewhere (except for orbital part of superior frontal gyrus, $p>0.05$, Mann-Whitney U test, two-sided, FWER corrected). **b**, Spatial distribution of variation in sliding-windowed PSDs during 5-min resting (left) and 60-min wakefulness (right), respectively. In the box plot **a**, the box extends from the first quartile to the third quartile of the data, with a line at the median. The whiskers extend from the box to the extreme data points, and outliers are shown as fliers. Asterisks show significance level of $*p\leq0.05$, $**p\leq0.01$ and $***p\leq0.001$.

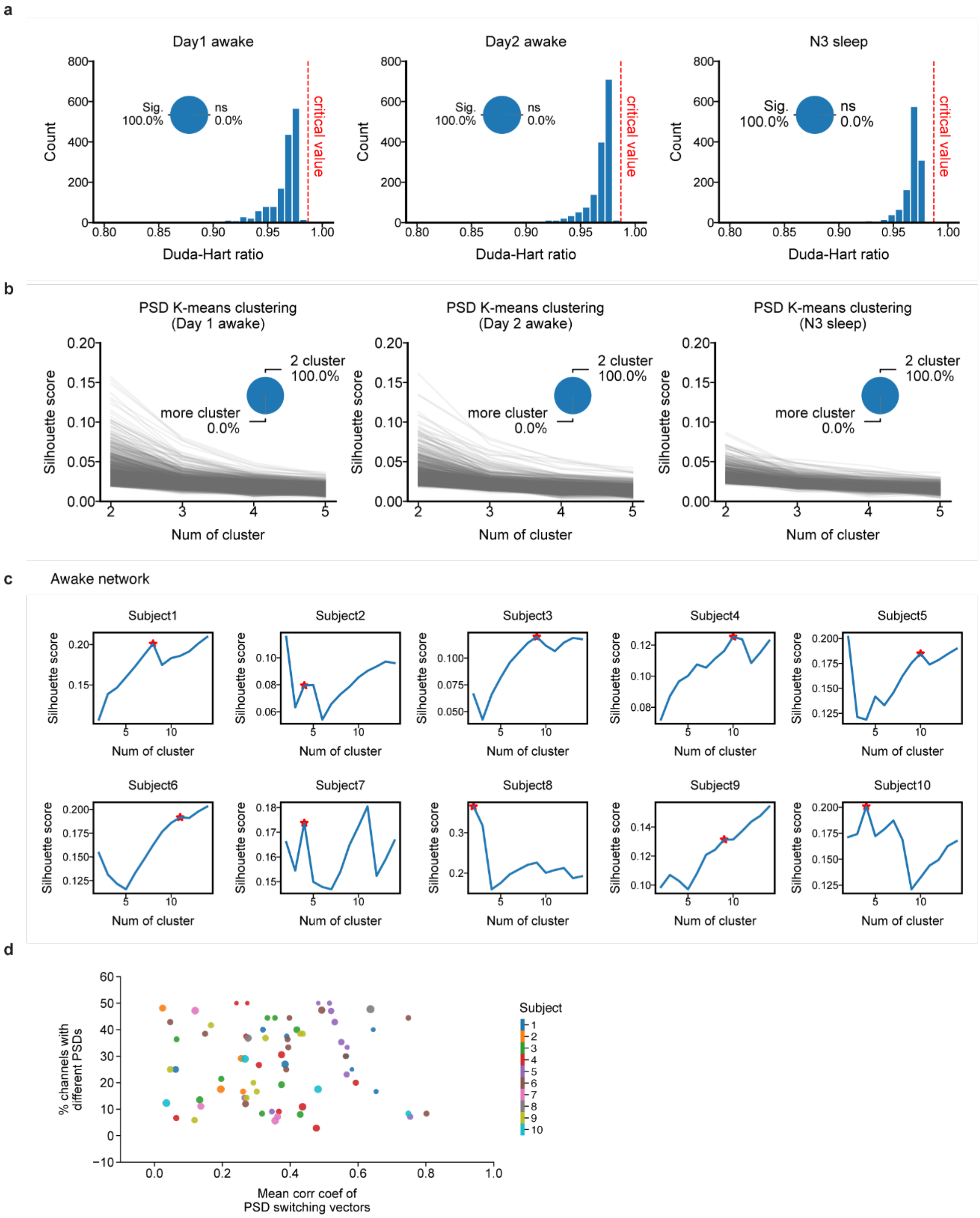

**Fig. S2 Number of clusters in sliding-windowed PSD profiles and functional networks.** a, Distribution of Duda-Hart ratio for each individual channel in all subjects during the 60-min awake period on day 1 (Left, n=1513), day 2 (Middle, n=1500) and N3 sleep state (Right, n=1189), as shown in Fig. 3, Fig. 6 and Fig. 7. The red dashed line indicates the critical value for two clusters. Across all subjects ($p<=0.001$ in all examined channels, FWER corrected). b, Silhouette scores for each number of clusters in PSD profile K-means clustering for individual channels. Each line represents the results of one channel during the 60-min awake period on day 1 (Left), day 2 (Middle) and N3 sleep state (Right). **c**, Silhouette scores for each number of clusters in network characterization from day 1 wakefulness (as shown in **Fig. 3**). The red star in each subplot indicates the selected optimal number of clusters within each subject. **d**, Mean correlation of PSD switching vectors within each network versus percentage of channels showing different PSDs (as shown in **Fig. 3**). Color indicates individual subject; size indicates the total number of channels within each network.

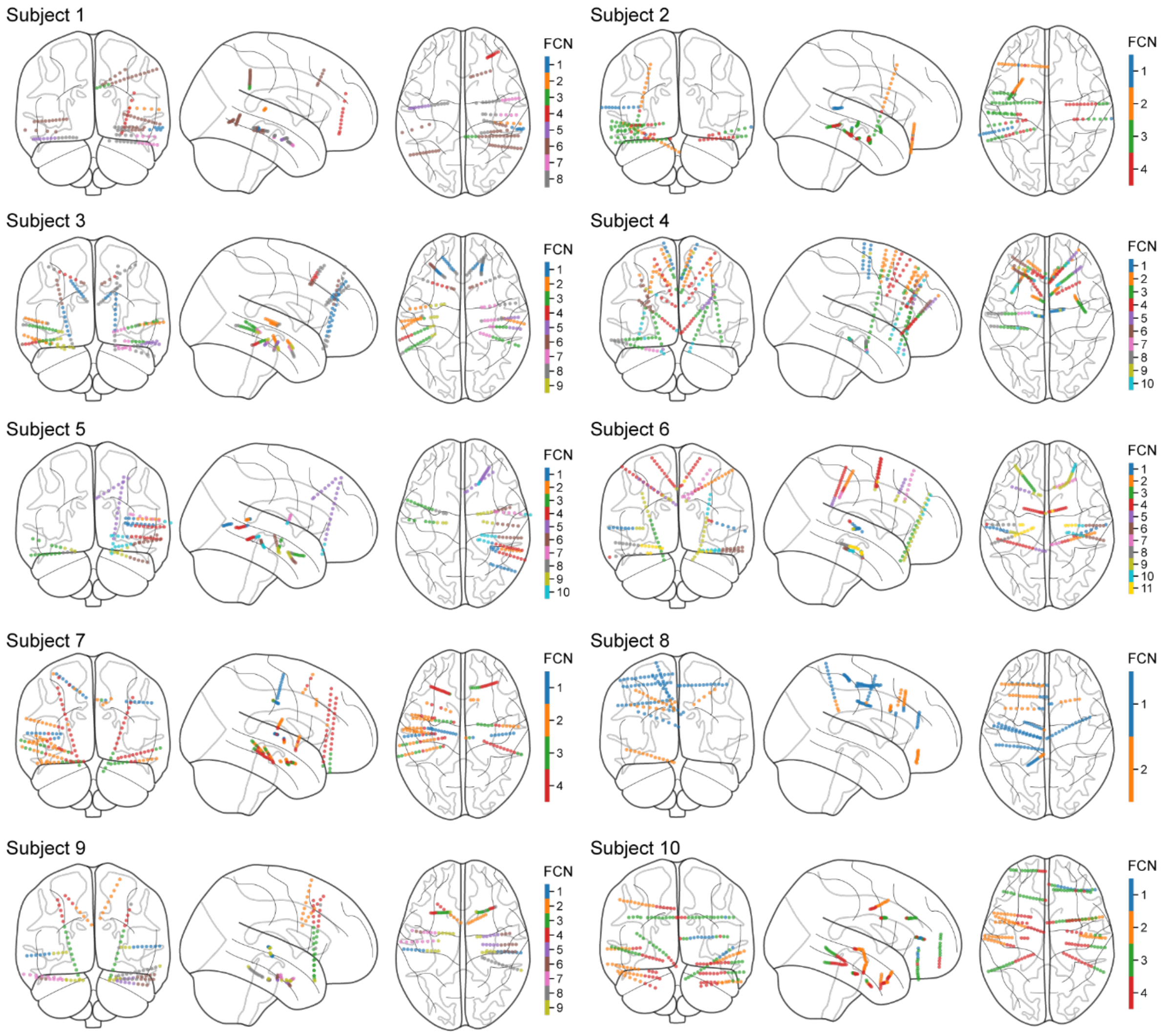

**Fig. S3 Networks in individual subjects characterized from day 1 wakefulness.** Subject-specific networks were characterized using the hierarchical clustering based on the correlation distance between channel's PSD switching vectors (as shown in **Fig. 3**). The number of networks varied across patients, depending on the electrode implant configuration for clinical evaluation purposes. Networks were found to span long distances and/or multiple brain areas, with bilateral networks observed in all subjects.

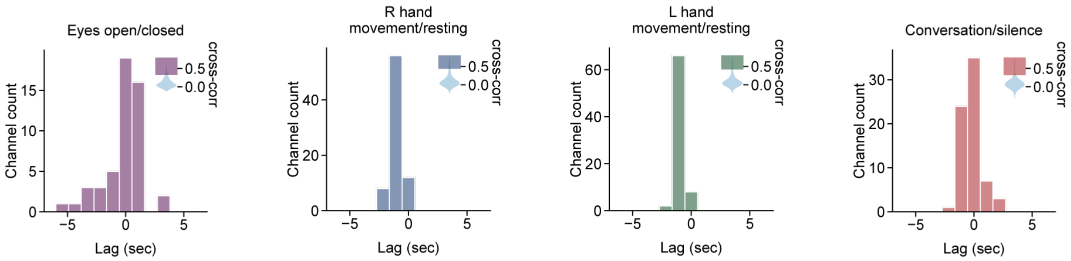

**Fig. S4 Optimal time lag between PSD switching and behavioral annotation.** Distributions of optimal lag that maximizes the cross-correlation between PSD switching vectors and behavioral state transition annotations for eyes open/closed, right-/left-hand movement/resting and conversation/silence, from left to right, respectively (as shown in **Fig. 5**). The violin plot shows the distribution of maximal correlation coefficients of all channels across subjects, and the histogram shows the distribution of optimal lag in channels whose correlation coefficients above the 95th quantiles (n=50, n=76, n=76, n=70 channels).

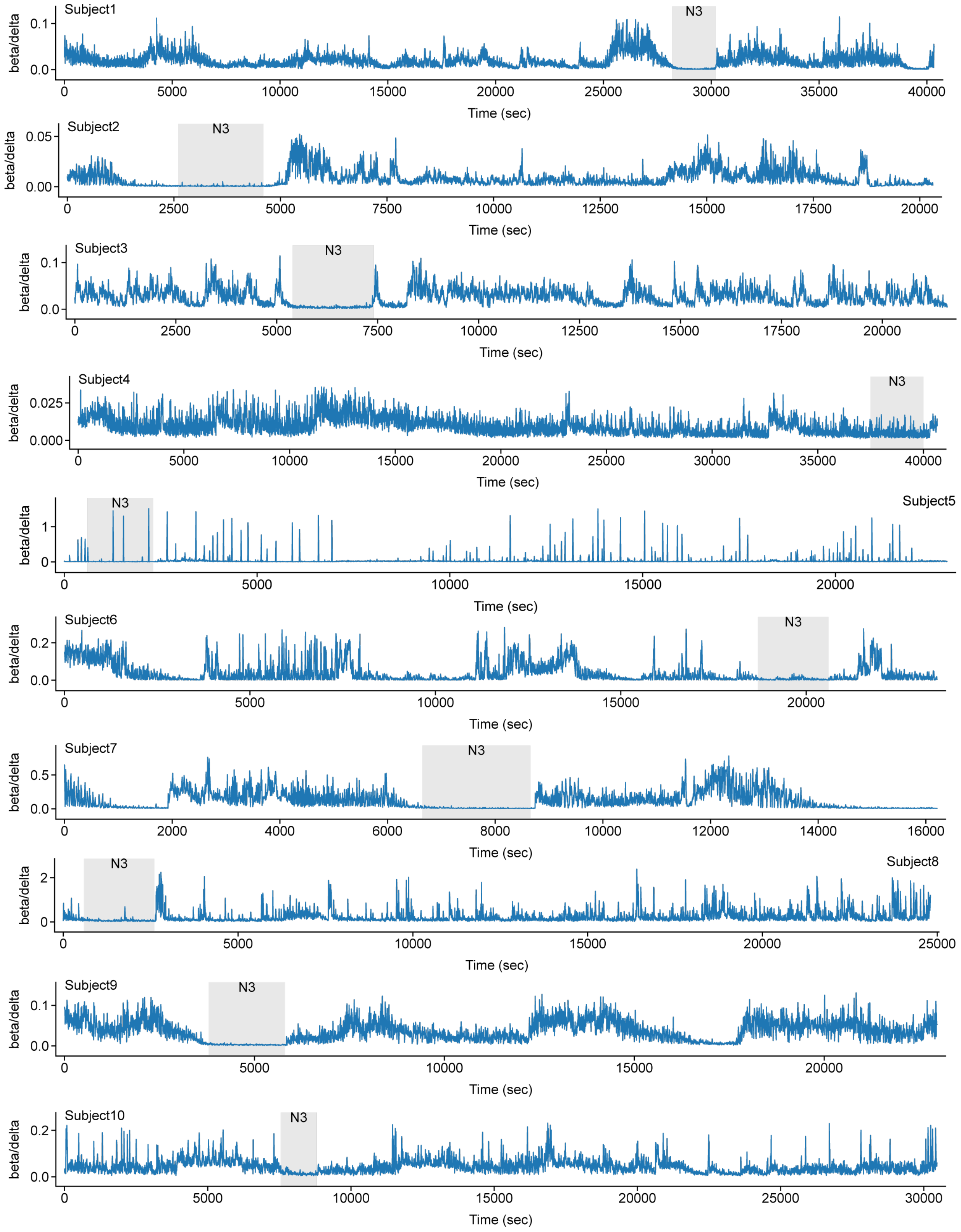

**Fig. S5 Full night sleep data and selected N3 sleep state Beta/delta power ratio across full night sleep for each subject.** Shaded areas indicate the selected N3 sleep state (as shown in **Fig. 6** and **Fig.** 7). Subject 4 and 5 were excluded from the network comparative analysis due to undetermined N3 sleep.

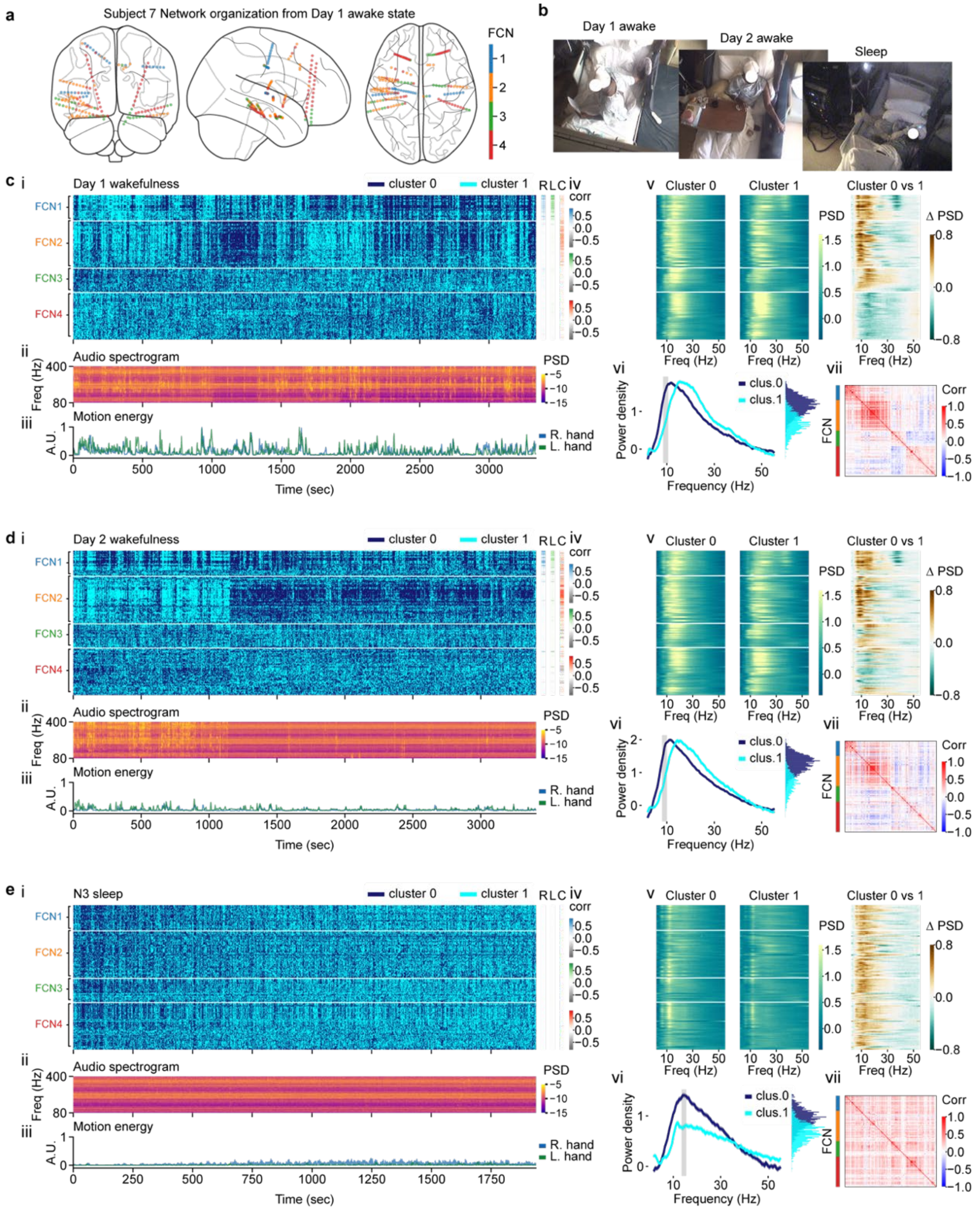

**Fig. S6 Comparing network dynamic results across days and sleep state (subject 7, as shown in Fig. 6). a**, Network organization characterized from day 1 wakefulness. **b**, Screenshots of monitoring video from day 1 awake, day 2 awake and N3 sleep period. **c**, (i): PSD switching vectors of individual channels during day 1 wakefulness, listed by networks. (ii): Audio spectrogram from simultaneous audio-visual recordings. (iii): Motion energy of DLC-labeled left/right-hand positions from simultaneous audio-visual recordings. (iv): Correlation coefficients of individual channels in response to each category of behavioral changes (right (R) and left (L) hand motion energy, and audio power (C)). (v): Mean PSD profiles of clusters 0 (left) and 1 (middle) from K-means clustering in individual channels, and their differences (right), listed by networks shown in subplot **c** (i). (vi): Mean PSD profiles (mean±1.96 SEM) of each cluster from an example channel located in left superior temporal pole, along with distribution of power averaged between 8.5-10.5 Hz (p≤1x $\mathbf{10^{-6}}$, Mann-Whitney U test, two-sided), color-coded by the K-means clusters. (vii): Correlation matrix of PSD switching vectors of individual channels. **d**, Results from day 2 wakefulness, following the same configuration as shown in subplot **c**. **e**, Results from N3 sleep state, following the same configuration as shown in subplot **c**.

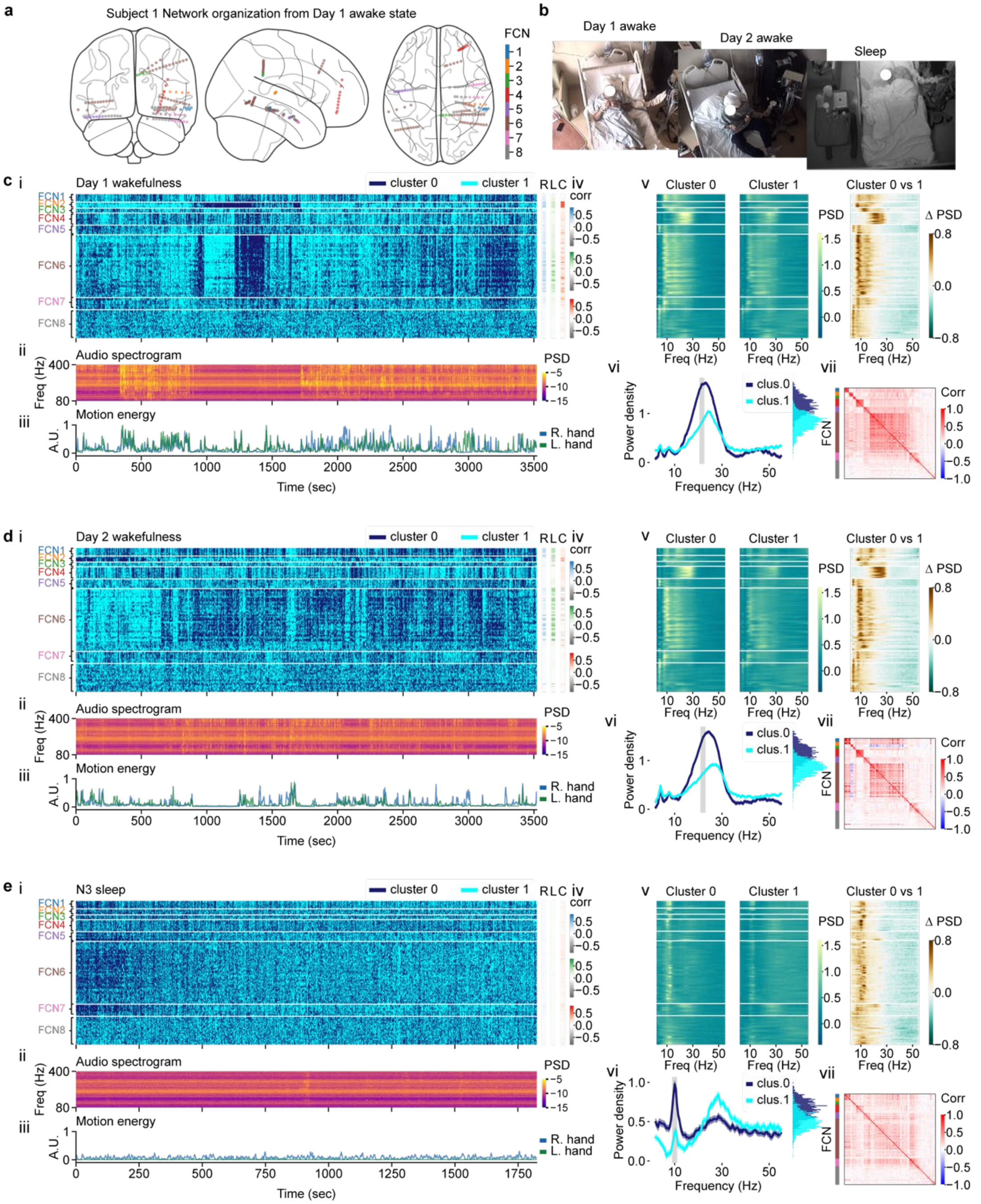

**Fig. S7 Comparing network dynamic results across days and sleep state (subject 1, as shown in Fig. 6).** **a**, Network organization characterized from day 1 wakefulness. **b**, Screenshots of monitoring video from day 1 awake, day 2 awake and N3 sleep period. **c**, (i): PSD switching vectors of individual channels during day 1 wakefulness, listed by networks. (ii): Audio spectrogram from simultaneous audio-visual recordings. (iii): Motion energy of DLC-labeled left/right-hand positions from simultaneous audio-visual recordings. (iv): Correlation coefficients of individual channels in response to each category of behavioral changes (right (R) and left (L) hand motion energy, and audio power (C)). (v): Mean PSD profiles of clusters 0 (left) and 1 (middle) from K-means clustering in individual channels, and their differences (right), listed by networks shown in subplot **c** (i). (vi): Mean PSD profiles (mean±1.96 SEM) of each cluster from a channel located in right middle frontal gyrus (orbital part), along with distribution of power averaged between 20.5-22.5 Hz ($p \leq 1\text{x } \mathbf{10^{-6}}$, Mann-Whitney U test, two-sided), color-coded by the K-means clusters. (vii): Correlation matrix of PSD switching vectors of individual channels. **d**, Results from day 2 wakefulness, following the same configuration as shown in subplot **c**. **e**, Results from N3 sleep state, following the same configuration as shown in subplot **c**.

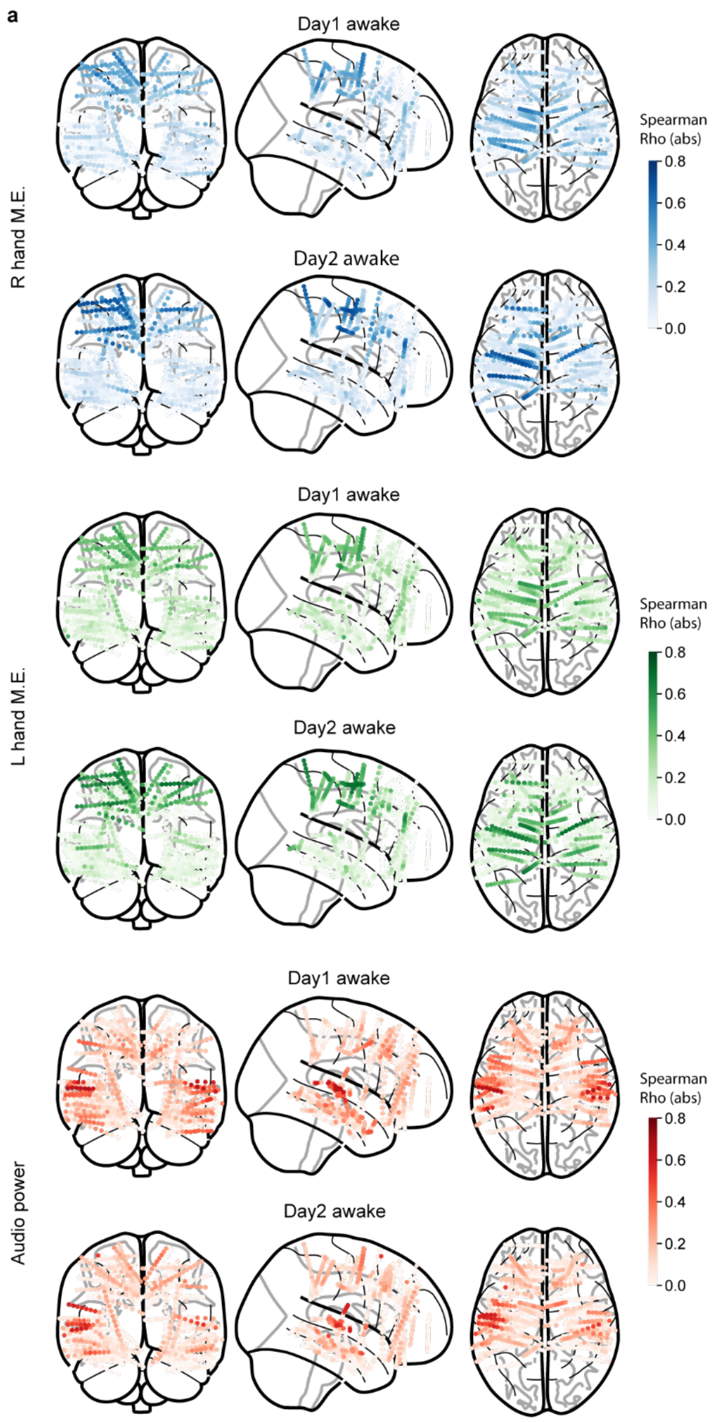

**Fig. S8 Correlation between PSD switching vectors to extracted left/right hand motion energy and audio power from simultaneous audio-visual recordings during wakefulness on day 1 and 2.** Spatial mapping of individual channel's correlation coefficients on day1 (top) and day 2 (bottom) onto brain plots for right hand, left hand and audio power, respectively (as shown in **Fig. 6** and **Fig.** 7).

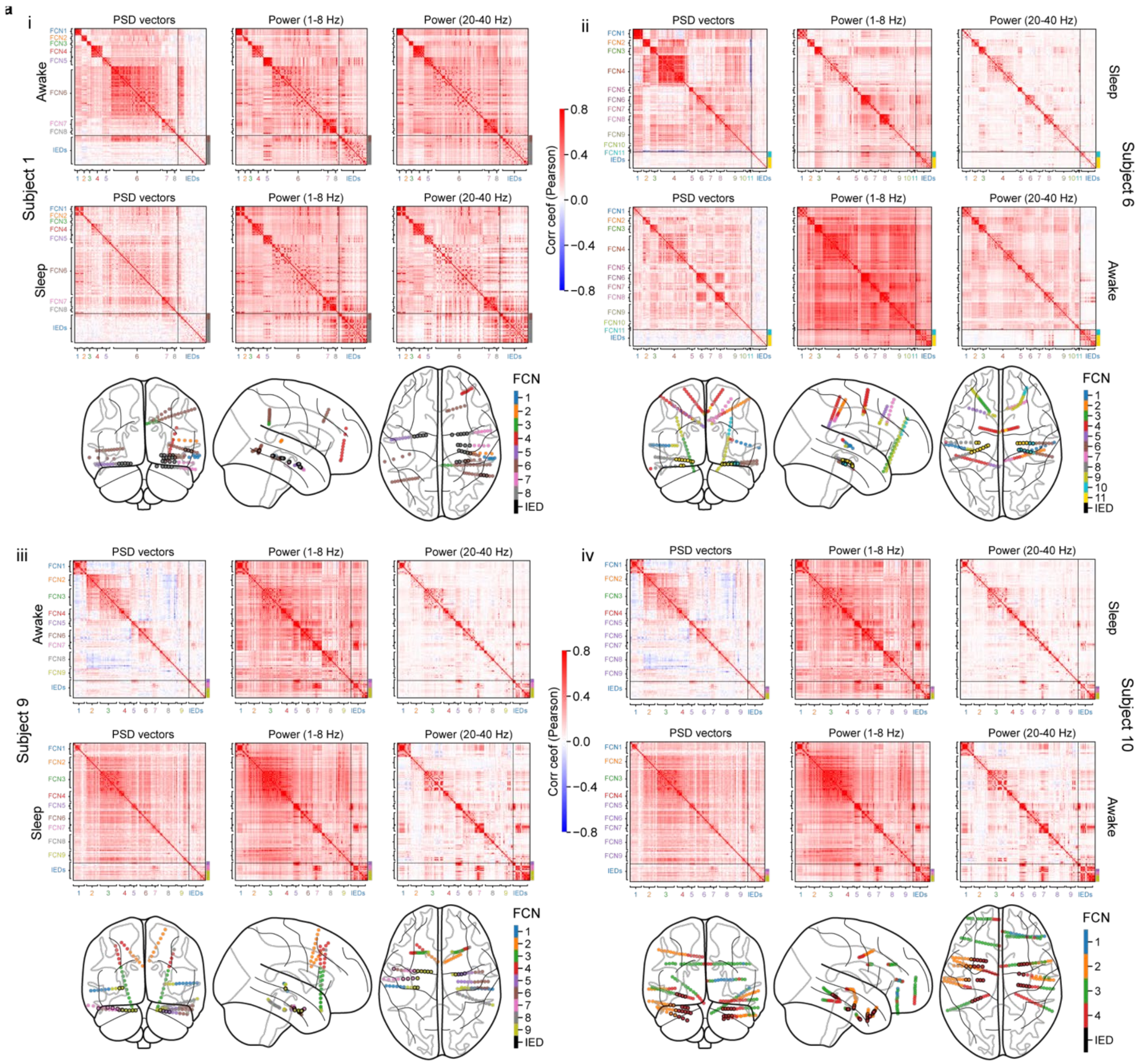

**Fig. S9 Correlation map and inter-network distance changes between normal channels and channels with frequent interictal epileptiform discharges (IED) during sleep. a**, Correlation maps based on PSD switching vectors (left), low frequency (1-8 Hz, mid) and high frequency (20-40 Hz, right) power during day 1 wakefulness (top) and N3 sleep state (bottom) from example subject 1 (i). Normal channels are arranged according to their assigned networks identified during day 1 wakefulness, IED channels (as shown in **Fig.** 7) are positioned at the bottom of the map. Correlation maps from subject 6 (ii), subject 9 (iii) and subject 10 (iv) are presented with similar configurations as shown in subplot **a** (i).

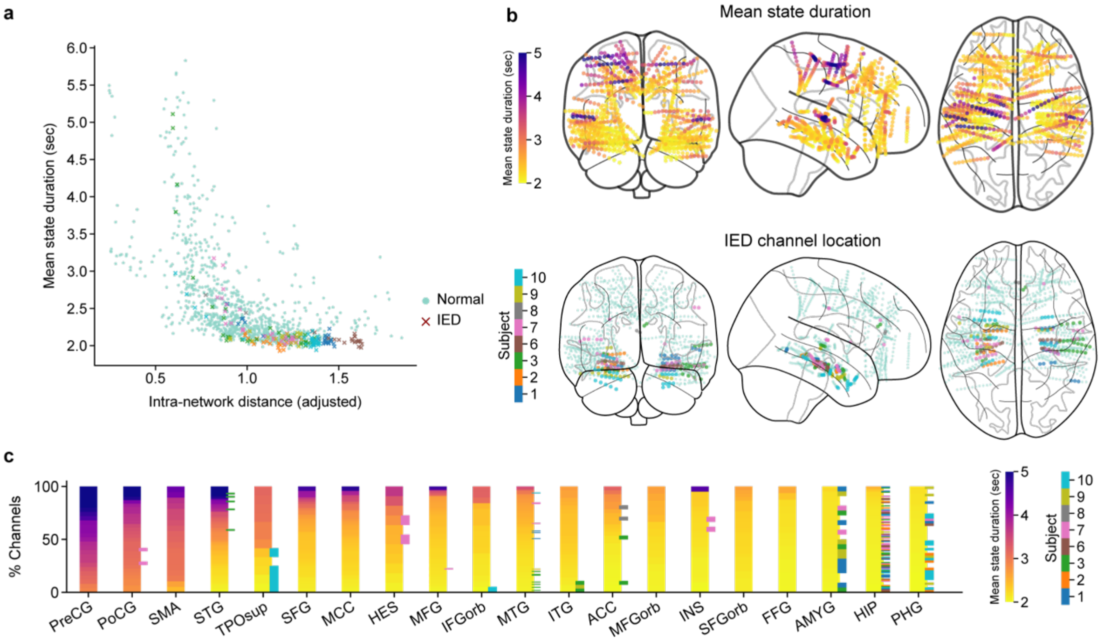

**Fig. S10 Mean state duration brain mapping and IED channel distributions. a**, Mean state duration vs intra-network distance (awake, adjusted); dots indicate normal channels, crosses indicate IED channels (as shown in **Fig.** 7) and color-coded by subjects. **b**, Top: Individual channel's mean state duration onto brain plots. Bottom: location of IED channels on to brain plots, color-coded by subjects. **c**, Distribution of mean state duration mapped onto brain regions of interest; regions are listed from the longest to the shortest region-averaged state duration from left to right. Horizontal lines indicate IED channels, with colors indicate corresponding subjects. IED channels can exhibit both long state durations and short state durations (fast-switching), and are found in regions with both long mean state durations and short state durations.
